# Extended discrete gene regulatory network model for the *Arabidopsis thaliana* root-hair cell fate

**DOI:** 10.1101/2023.11.15.567304

**Authors:** Aarón Castillo-Jiménez, Adriana Garay-Arroyo, M. de La Paz Sánchez, Berenice García-Ponce, Juan Carlos Martínez-García, Elena R. Álvarez-Buylla

## Abstract

The differentiation of the two cell types of the root epidermis, atrichoblasts, which give rise to hair cells, and atrichoblasts, which do not develop as hair cells, is determined by a complex regulatory network of transcriptional factors and hormones that act in concert in space and time to define a characteristic pattern of rows of hair cells and non-hair cells interspersed with each other throughout the root epidermis of *Arabidopsis thaliana*. Previous models have defined a minimal regulatory network that recovers the Wild Type phenotype and some mutants but fails to recover most of the mutant phenotypes, thus limiting its ability to spread. In this work, we propose a diffusion-coupled regulatory genetic network or meta-Gene Regulatory Network model extended to the model previously published by our research group, to describe the patterns of organization of the epidermis of the root epidermis of *Arabidopsis thaliana*. This network fully or partially recovers loss-of-function mutants of the identity regulators of the epidermal cell types considered within the model. Not only that, this new extended model is able to describe in quantitative terms the distribution of trichoblasts and atrichoblasts defined at each cellular position with respect to the cortex cells so that it is possible to compare the proportions of each cell type at those positions with that reported in each of the mutants analyzed. In addition, the proposed model allows us to explore the importance of the diffusion processes that are part of the lateral inhibition mechanism underlying the network dynamics and their relative importance in determining the pattern in the Wild Type phenotype and the different mutants.

## Introduction

The *Arabidopsis thaliana* root epidermis is an ideal complex biological system to explore the generic mechanisms that underlie both cell fate and pattern formation in plant development. This plant structure consists of a mono-layer of two types of cells: the trichoblasts (root-hair cells) that could form root hairs, and the atrichoblasts (non-hair cells) that cannot form hairs. Epidermal cells between two underlying cortical cells (H-position) adopt the root-hair cell fate and cells adjacent to only one underlying cortical cell (N-position) adopt the non-hair cell fate. Thus, a specific organization pattern of the root epidermis is established: root hairs are arranged in columns and these hairs columns are interspersed by columns containing only non-hairs cells [24, 34, 37, 49, 105]. Transverse divisions of the root meristem originate new epidermal cells, that differentiate progressively while retaining the previously established position-dependent pattern in earlier stages of their development [14, 35, 37, 51, 68, 102]. Thanks to both the presence of few cell types in the *Arabidopsis thaliana* root epidermis and the ease of dynamically tracking them during their specification and differentiation, it is possible to follow a formal model-based systems biology approach to study root epidermal differentiation and systemically elucidate the underlying generic regulatory mechanisms [50, 52, 75, 76].

The molecular regulatory mechanisms that regulate both the process of differentiation and the development of the *Arabidopsis thaliana* root epidermis have been extensively studied and reviewed: In cells in N-position, WEREWOLF (WER), GLABRA3 / ENHANCER OF GLABRA 3 (GL3 / EGL3) and TRANSPARENT TESTA GLABRA 1 (TTG1) form a MBW (MYB, bHLH, and WDR) complex or *transcription activation complex* [49]. This complex promotes the transcription of GLABRA2 (GL2) and CAPRICE (CPC). GL2 then begins to accumulate and inhibit the activity of root-hair-promoting genes. CPC begins to mobilize and accumulate in cells in the H-position where it competes with WER and forms a complex with TTG1 and GL3 / EGL3 or inhibitory complex which causes a decrease in the expression of GL2 allowing for transcription of root-hair-promoting genes and root-hair differentiation. In parallel, in H-position cells the SCRAMBLED (SCM) activity is higher allowing the MBW complex activity to decrease by inhibiting WER [48, 49, 56, 101, 110, 113, 124]. In turn, WER, as part of the MBW complex in the N-cells, inhibits the activity of SCM while CPC and TRY promote it [66]. Altogether these regulatory mechanisms establish the robust pattern of spatial cellular organization in the root epidermis of *Arabidopsis thaliana* [11, 108].

Previous works have integrated the regulatory mechanism described above in gene regulatory network (GRN) models [9, 11, 56, 102]. Such networks have been profusely used to uncover the concerted activity of the transcriptional factors that participate in the determination of these cells as well as in the spatiotemporal dynamics behind the characteristic pattern of distribution of trichoblasts and atrichoblasts in *Arabidopsis thaliana* root [9, 11, 85, 103, 114]. Moreover, it has been established that the characterized networks give rise to the dynamic structural mechanisms that determine the observed organizational pattern. Which is to say: the existence of a mechanism of lateral inhibition through the accumulation of the MBW complex that regulates the expression of diffusible molecules (CPC, TRY, and ETC1) that prevent adjacent cells from adopting the same fate through inhibitory complex activity; positive feedback loops on WER activity (through the MYB23 activity); as well as among the positional signals coming from the cortex through the SCM and JACKDAW (JKD) activity have been described [59, 100, 108, 140].

The referred network-based mathematical models, influenced by the pioneering works of Turing [129] and Meinhardt [47, 83], have suggested the existence of generic patterning mechanisms, such as self-regulation and long-range inhibition [100]. A dynamic activator-inhibitor mechanism has been suggested as the main dynamic process to explain the spatial pattern of the root epidermis. This mechanism generates concentration differences between a self-promoting activator (the MBW complex) and an inhibitor with high intercellular mobility (the IC). Along with this key pattern-forming regulatory motif, positive and negative feedback loops between WER/MYB23 and CPC as well as among the positional signals coming from the cortex have been described to date [100, 108, 140].

In previous works [9–11, 13] our research group proposed a spatial network model (so-called meta-GRN model) consisting of a grid of 20 rows and 20 columns, in which each cell corresponds to a cell of the epidermis of the root of *Arabidopsis thaliana* and contains a GRN conformed by the genes WER, GL3, EGL3, TTG1, GL2, CPC, TRY, ETC1 and SCM. The proteins GL3, EGL3, and CPC can move passively between cells and can affect the expression of the genes of the network. In this model, the expected expression profiles for the cells corresponding to trichoblasts and those corresponding to atrichoblasts were recovered *in silico* for WT and some mutants [11]. We have also explored structural and functional properties of the networks, such as the presence of functional modules, and we have suggested a key role of the proposed feedback loops in the robustness of the spatial patterns of cellular organization in the root epidermis (See [9]). Since recent evidence points to the existence of additional regulators to those previously incorporated in the dynamic models. In our current study, we propose a discrete spatial GRN model that integrates the previously reported one with new transcriptional regulators and hormones that have been shown to be important in the root hair patterning. Our results show that the GRN model that we expose here recovers in the spatial patterning of the root epidermis in WT and various mutant backgrounds and hormonal conditions.

## Materials and Methods

### Discrete mathematical modeling

In order to uncover the systems-based mechanisms that underlie cell fate decisions in our case-of-study, we first proceed to update a discrete network model that we have previously developed [9–11, 13]. For this, we proceed via an extensive review of available curated literature on the field. Thus, we identified new regulatory interactions between the existing agents and we also incorporated some new regulators.

For simplicity, the state of the nodes in the GRN was updated synchronously. In the single-cell GRN, as was previously demonstrated in [11], the updating scheme does not seem to affect attractors when these are fixed-point types, yet this issue has not been explored in coupled GRN models.

In the GRN model, nodes of the network correspond to genes, and each node’s state depends on that of other nodes, its regulators. The level of expression of a given gene is represented by a discrete variable that takes its value in the discrete set *{*0, 1, 2*}*, where 0 denotes the low level of expression, 1 denotes the medium level of expression, and 2 denotes the high level of expression, *i.e. gɛ {*0, 1, 2*}* and it depends on the level of expression of other components of the network *g*_1_*, g*_2_*, …, g_N_*. The state of every gene *g* therefore changes according to:

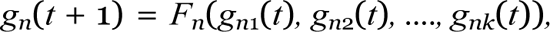

where *g_n_*_1_*, g_n_*_2_*, …, g_nk_*, are the regulators and *F_n_* is a logical regulatory function; *t* stands for the discrete time independent variable *t* = 0, 1, 2, ….

Given the logical rules, it is possible to follow the dynamics of the network for any initial configuration of the node’s expression states. In order to couple the GRNs in a compartmentalized domain, we considered a discrete lattice of *n × n* elements in which each element (*i, j*) represents a cell with a GRN and all cells in the lattice bear the same GRN. Thus, modeled lattices are a simplified representation of root epidermal sections.

We must point out that cells in the lattice have exactly four neighbors and there is no difference in permeability between them.

According to experimental data, some proteins codified by elements of the network (CPC, GL3, and EGL3) move to neighbor cells and affect gene expression in a non-cell-autonomous fashion giving rise to a network of coupled networks (herein meta-GRN). Although empirical evidence supporting cell-to-cell motion rather than aplopastic transport only exists for CPC, all available data are congruent with the assumption that all mobile elements of the GRN move in a cell-to-cell manner. In the spatial model we therefore allowed these elements to move among neighboring cells following the process described by the equation:

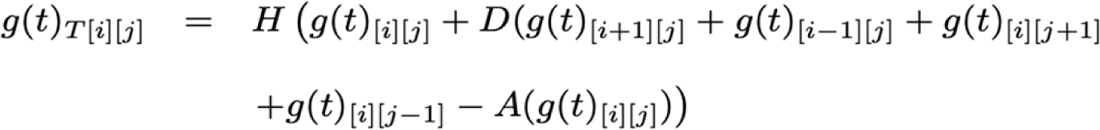

where:

- *g*(*t*)*_T_* _[*i*][*j*]_ is the total amount of protein *g* in cell(*i, j*).
- *D* is a continuous variable that determines the proportion of *g* that can move from any cell to neighboring ones and is correlated to the diffusion rate of *g*.
- *H* is a step function that converts the continuous values corresponding to the amount of *g* diffused into each cell into a discrete variable that may attain the levels of expression characterized by *{*0, 1, 2*}*.

For modeling the root system two borders were identified and two were kept at zero-flux, simulating a root-like cylinder. For this, diffusion was allowed between the cells at the left end of the grid and the cells at the right end. In the cells found at the upper or lower ends of the cell grid, *A* adopts a value of 3 because these cells only have three neighboring cells towards which they can diffuse. In the rest of the cells, *A* has a value of 4 because they have 4 neighboring cells towards which the mobile elements can diffuse. Then, *g*(*t*)*_T_* _[*i*][*j*]_ was considered in order to estimate the *g*(*t*)_[*i*][*j*]_ value, which was then used to update the logical rules, according to the next equation:

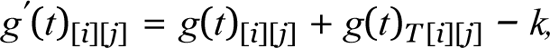

where *k* is a specific degradation constant for each moving element. Thus, *g^′^*(*t*)_[*i*][*j*]_ was taken as an input to evaluate the logical rules and obtain *g*(*t* + 1)_[*i*][*j*]_. In brief, each cell’s GRN is initialized with a random gene activation profile except for MYB23, TTG1, and WRKY75 which have expression profiles of 0, 1, and 1 respectively (see the reasons for this later in the results section). Then, three steps are sequentially repeated until the whole lattice reaches a steady state (See Figure 1 for the corresponding schematic sequence). After 1,000 iterations, cells that express the GL2 gene are considered atricoblasts, while those that do not express it are considered trichoblasts. Thus, a data matrix is formed in which the cells that correspond to trichoblasts have a value of 1 and those that correspond to atricoblasts have a value of 0. With this data matrix, the spatial pattern of distribution of trichoblasts and atricoblasts is plotted (See Figure 1). For each simulation In order to test the model, we also simulated networks that correspond to reported mutants. The loss and gain of function simulations were done by fixing the expression value of the altered gene to 0 or 2, respectively, and the spatial pattern obtained at the end of the algorithm was reported for each one and compared with the phenotypes reported in the literature for each mutant that has been taken into account. For each simulation the total number of trichoblasts and atricoblasts in each position of the grid was counted. To have a statistically significant measurement, a total of 10,000 simulations were performed and the average percentage of both cell types was calculated for each considered phenotype. We must point out that the random seed was the same for both wild-type and mutant simulations so that results were comparable. The resulting graphics were elaborated in Matlab and the corresponding computer programs were written in C language (codes available upon request).

**Figure 1.**
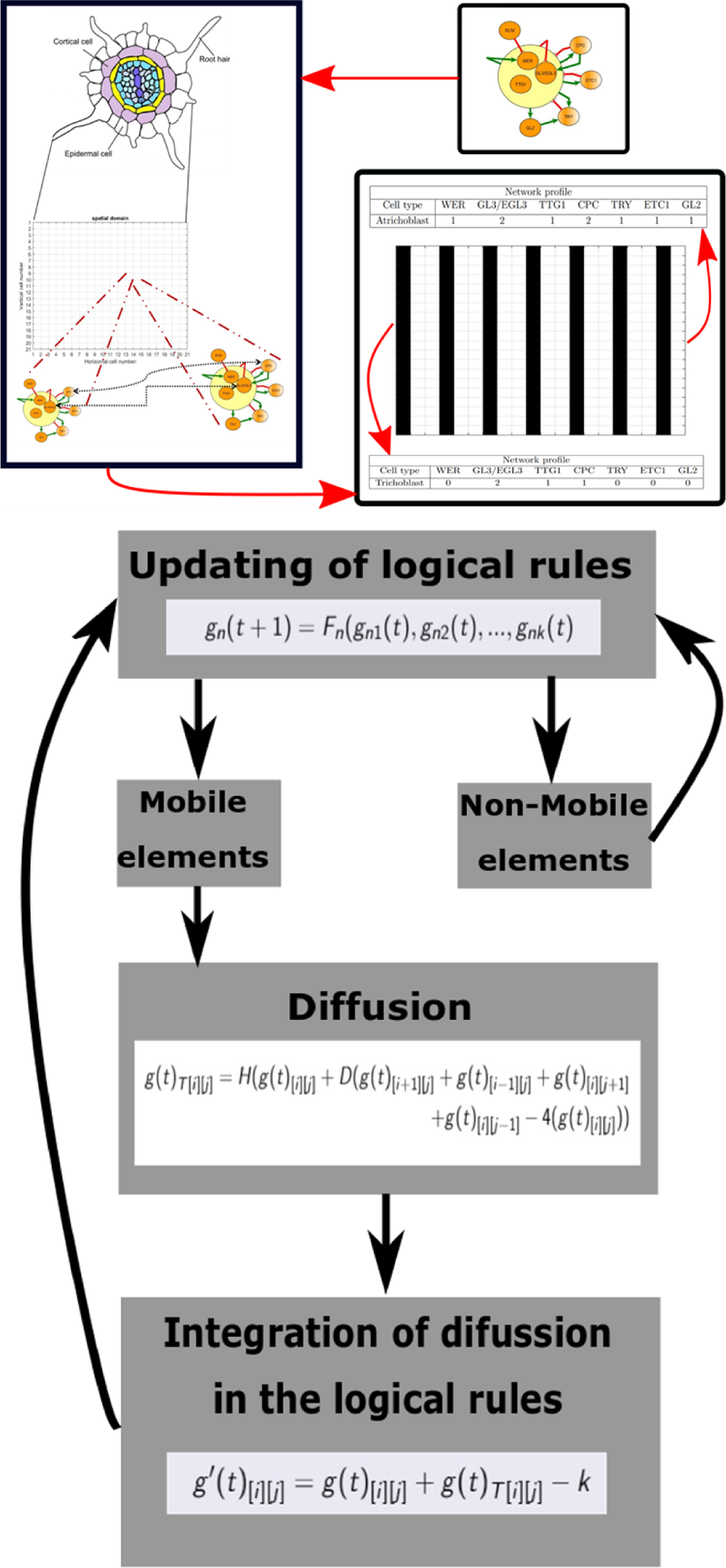
Algorithmand simulation scheme.

## Results

### Updated epidermal cell subdifferentiation meta-GRN model

To extend our previous meta-GRN model [9, 11,13], we extensively reviewed the available experimental information until 2021. Then, we decided to include the genetic components that were well characterized at the functional level in wild-type and various mutant conditions, particularly, documented functional feedback regulatory loops in the network models that have been proposed related to hormonal signaling pathways of Auxins, Cytokinin, and Ethylene (See Table SP1 in Supplementary Information). Thus, after an exhaustive review of the experimental information, new genes were integrated into the GRN. These are the following ones: atMYB23 (MYB23), TRANSPARENT TESTA GLABRA 2 (TTG2), WRKY75, ZINC FINGER PROTEIN5 (ZFP5) and JACKDAW (JKD). We summarize in Figure 2 the GRN topology and their interactions with all the new and previous nodes and interactions used for the meta-GRN model. All detailed information about interactions can be found in Supplementary Information.

**Figure 2.**
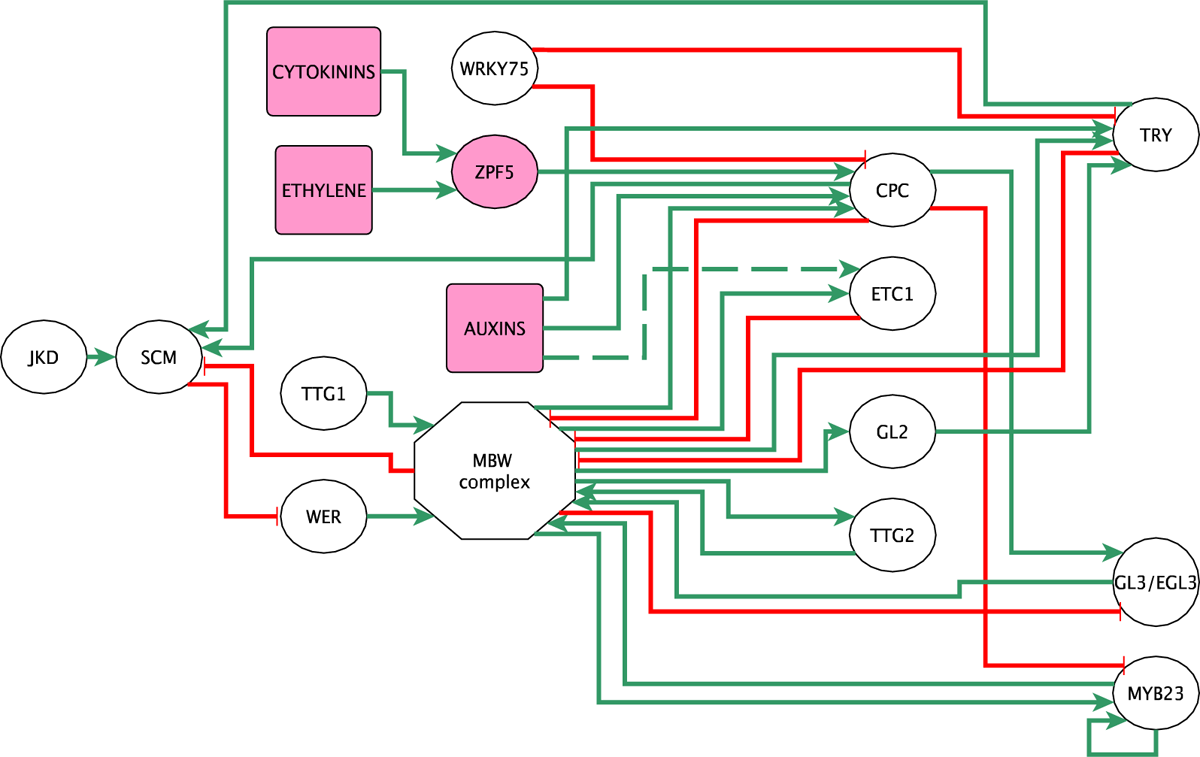
GRN topology. The red lines indicate repressive transcriptional interactions while the green lines indicate activating transcriptional interactions. The two nodes in pink are nodes that refer to hormones, the two with two colors are nodes that participate in the integration of hormonal signals. The round nodes refer to genes, the square ones to hormones and the hexagonal node represents a protein complex.

In what follows, the elements that are part of the MBW complex, the inactivating complex, and the canonical elements of the network that describe the sub-differentiation of the epidermis as well as the newly incorporated regulators are described one by one. In each case, the experimental evidence supporting the involved regulatory interactions is also specified.

WER is an important regulator of root hair identity and a central component of this regulatory network: On the one hand, it regulates the activity of GL2 in the determination of non-hair cells and, on the other hand, it regulates the activity of CPC to determine the identity of the root hair cells [63, 72, 99]. WER encodes an R2R3 MYB transcription factor [72, 120] and its expression takes place only in cells in N-position, whereas it is suppressed in H-position cells [65, 68, 72]. In the cells in N-position WER physically binds to the bHLH transcription factors GL3 and EGL3 and with the WD40 protein TTG1 to form an activator complex or the MWB complex [16, 92], whose function is to activate the GL2 gene in cells in N-position by direct binding of WER to the promoter of GL2 [63], which inhibits the generation of root hair and thus makes cells differentiate into N-position cells or Non-hair cells [16, 63, 72, 92, 135, 145].

WER activity is negatively regulated by SCM since it has been observed that in the *scm-2* mutant an increase in WER transcription is observed, while the over expression of SCM in the 35S::SCM and WER::SCM mutants decreases the expression of WER [65]. On the other hand, SCM preferentially accumulates in cells in position H and since in the *scm-2* mutant, an alteration of the pattern of organization with respect to the position of cortex cells is observed [68]. This suggests that SCM functions as a positional regulation signal on WER expression thus inhibiting its activity in H-position cells thus allowing the identity of trichoblasts [68]. Mutations in WER give rise to roots with hairy phenotypes. In the *wer-1* mutant the expression of GL2 is almost completely reduced so hairs are observed in all the cells of the epidermis [73]. WER is negatively regulated by SCM since it has been observed that in the *scm-2* mutant an increase in WER transcription is observed, while the overexpression of SCM in the 35S::SCM and WER::SCM mutants decreases the expression of WER [65]. On the other hand, SCM preferentially accumulates in cells in position H and since in the *scm-2* mutant, an alteration of the pattern of organization with respect to the position of cortex cells is observed [68]. This suggests that SCM functions as a positional regulation signal on WER expression thus inhibiting its activity in H-position cells thus allowing the identity of trichoblasts. It has been suggested in the literature, that WER presents a direct or indirect positive feedback interaction since in *cpc-1* mutants the transcription of WER increases and its ectopic expression is observed in some H-position cells [73]. The possibility that WER regulates its activity itself is not conclusive given that in the *wer-1* mutant there are no changes in the activity of the WER promoter, so it is not possible to establish that WER regulates itself [59]. However, it has been shown that the MBW complex positively regulates MYB23 whose protein is functionally similar to WER and MYB23 directly inactivates SCM, which allows WER activation. Thus it has been suggested that MYB23 has indirect positive feedback on WER activity, but this interaction has not been analyzed in previous GRN models [59, 66, 108]. In our previous theoretical works, we proposed the existence of a positive feedback mechanism in the formation of the MBW complex [1, 11], but the nature of this interaction had not been attributed to regulation through MYB23. MYB23 not only works in a manner analogous to WER in the formation of the AC but also its expression is preferably in the cells in N position [59, 109].

MYB23 encodes for a protein of the MYB family. The MYB23 protein is functionally similar to the WER [72]; MYB23 physically interacts with GL3/EGL3 and TTG1 to form a complex that is functionally similar to the AC [133]. The expression of MYB23 can be regulated by the WER promoter so that in the mutant line WER::MYMB23 *wer-1* it is possible to recover the normal expression of GL2 that is regulated by WER on the MBW complex. It is also observed that the levels of MYB23 RNA in the mutant line *wer-1* are reduced by 19% while the expression of the reporter PMYB23:GUS is almost undetectable, suggesting that WER induces the expression of MYB23 [59].

The WER and MYB23 protein physically interacts with the MYB23 promoter since it is observed that the WER protein binds in 4 sites in the promoter region of MYB23 [59]. MYB23 is also regulated by TTG1, CPC, GL3 and EGL3 genes. In the mutant lines *ttg1-13* and *gl3-1 egl3-1* a decrease in the accumulation of MYB23 RNA is observed, while a decrease in the accumulation of PMYB23::GUS is observed maintaining the pattern of the cells in N-position. In the case of the mutant line *cpc-1* an increase in the expression levels of the MYB23 transcripts of 56% with respect to the WT is observed, while in the P35S:CPC overexpression line no activity of the MYB23 promoter is observed when the reporter PMYB23:GUS is analyzed [59]. These results as a whole suggest that MYB23 is positively regulated by the WER-TTG1-GL3/EGL3 complex that promotes the identity of atrichoblasts, and negatively regulated by CPC that promotes the identity of trichoblasts. In the MYB23::MYB23-SRDX line that expresses a chimeric protein with a strong repressor activity the expression of CPC, GL2 and MYB23 is decreased, and an excessive root hair cell specification due to the expression of hairs in the N position. This indicates that MYB23 is a positive regulator of CPC, GL2, and its own expression [59]. With this information, we propose that the expression of MYB23 is regulated positively by elements of the MBW complex and by itself.

T R A N S P A R E N T T E S T A (TTG2) encodes for a WRKY transcription factor that is preferentially expressed in cell nuclei at the N-position [55,58]. Mutations in TTG2 have been shown to cause defects in trichome development but not in root hairs. In the *ttg2-1* mutant, no differences in root hair growth patterns are observed compared to the wild-type phenotype [58]. No GUS staining is seen in the *wer-1* mutants and the *gl3-7454 egl3-5712* double mutant lines in both ProTTG2:GUS and ProTTG2(2.0):GFPGUS lines. However, in the *cpc-2* mutant, ectopic staining is observed in some H cells, and in the *cpc-2 try-29760* double mutant, staining is observed in all H cells. These findings indicate that WER, GL3, and EGL3 have a positive regulatory effect on TTG2 expression in N cells, while CPC and TRY act as repressors of TTG2 expression in H cells [55]. It has been observed that there was no GUS staining in the *wer-1* and *gl3-7454 egl3-5712* double mutant and some GUS staining in H-position cells in the *cpc-2* single mutant and in all H-position cells in the *cpc-2 try-29760* double mutant using the ProTTG2:GUS line [55]. These results suggest that WER, GL3 and EGL3 positively regulate TTG2 expression in N-position cells and that CPC and TRY repress it in the H-position cells. Moreover, the CPC repression of TTG2 was confirmed only in leaves in with Pro35S:CPC transgenic plants [55]. In contrast, the ProTTTG2:TTG2:SRDX line, acting as a dominant repressor, exhibits distinct changes in gene expression compared to the wild type. In this line, the expression of GL2 is significantly reduced, while both CPC and endogenous TTG2 showequal reductions. Interestingly, GL3 and EGL3 exhibit significantly higher expression levels. However, the expression of TTG1 remains similar to that observed in the wild type. Furthermore, the analysis of ProGL2:GUS reporters reveals a decrease in expression, while ProTTG2:GFP:GUS shows a reduced expression that is not specific to the N-position cells. The ProCPC:GUS reporter does not show any expression in the epidermis. Conversely, the ProWER:GFP exhibits a strong fluorescence signal in N-position cells, but a weak signal in H-position cells. Based on the above findings, it can be inferred that TTG2 plays a regulatory role in the expression of CPC, GL2, and its own gene [55]. Finally, even when TTG2 is expressed in cells in the N position, no alterations are observed in the phenotype of the *ttg2-1* mutant, suggesting that other genes compensate for the loss of TTG2 function in this mutant [55]. In summary, the results indicate three key findings: firstly, WER, GL3, and EGL3 negatively regulate TTG2 expression, while CPC and TRY also have a negative regulatory effect. Secondly, the MBW complex and TTG1 positively regulate TTG2 expression. And lastly, TTG2 itself exhibits positive regulation of its own activity [55].

GLABRA3 (GL3) and ENHANCER OF GLABRA3 (EGL3) genes encode for bHLH transcription factors that share a close relationship [144]. The GL3 gene encodes a bHLH transcription factor, and EGL3 serves as its homolog, functioning redundantly with GL3 in Arabidopsis [17, 92] and have been shown to bind to similar promoter regions [88]. Although GL3 and EGL3 are expressed in H-position cells, their proteins accumulatein N-position cells through diffusion [17]. The mutant lines 35S:GL3 and 35S:EGL3, where GL3 and EGL3 are overexpressed, exhibit a reduced number of root hairs due to incorrect specification of H-position cells [17]. Although both cases display a similar phenotype, the impact is more pronounced in the 35S:EGL3 line [17]. When examining the *gl3-1*, *gl3-2*, *egl3-1*, and *egl3-2* mutants, it was observed that the lower region of the root exhibited a normal number and pattern of root hairs. However, in the upper region, *egl3-1* and *gl3-2* mutants displayed a slight increase in the number of hairs, while *gl3-1* and *egl3-2* mutants exhibited a moderate increase. This increase in the upper region was attributed to an abnormal specification of hairs in the N-position, specifically, an elevation in the occurrence of ectopic hairs, suggesting its importance in determining the identity of trichoblasts and atrichoblasts in the upper root region [16]. The analysis of GL3 and EGL3 double mutants, using combinations of the four mutant lines mentioned earlier (*i.e. gl3-1 egl3-1*, *gl3-1 egl3-2*, *gl3-2 egl3-1* and *gl3-2 egl3-2*), reveals an excessive production of root hairs and a decrease in the frequency of non-hairy cells along the root. This demonstrates that GL3 and EGL3 have partially redundant functions in determining the presence of non-hairy cells in the root. Collectively, these findings, along with the previously discussed results, provide further evidence for the partial redundancy of GL3 and EGL3 in this aspect of root development [16]. Interaction of GL3 and EGL3 with other elements of the genetic regulatory network has been demonstrated. In the double mutants *35S:GL3 wer-1* and *35S:EGL3 wer-1*, an excessive hair phenotype resembling *wer-1* is observed, indicating that the activity of GL3 and EGL3 is dependent on WER [16]. On the other hand, in the *gl3-1* mutant, there is a significant decrease in the expression of GL2, but the spatial arrangement of trichoblasts and atrichoblasts remains unaffected. In the *egl3-1* mutant, there are no changes in the expression of GL2. However, in the double mutant *gl3-1 egl3-1*, there is a notable decrease in GL2 expression, accompanied by an excessive abundance of root hairs [16]. These findings suggest that GL3 and EGL3 function redundantly to positively regulate the expression level of GL2. However, it does not affect the expression pattern of GL2. In contrast, when examining the *35S::GL3 gl2-1* and *35S::EGL3 gl2-1* double mutants, a hairy phenotype similar to that of the *gl2-1* mutant is observed. This similarity implies that both GL3 and EGL3 genes play a role in the determination of atrichoblasts through GL2 activity [16].

Regarding CPC activity, it has been observed that in the *gl3-1* and *egl3-1* mutants, as well as in the *gl3-1 egl3-1* double mutant, there is a reduction in the expression of the CPC::GUS reporter in the first case, no alteration in the second case, and almost complete elimination of expression in the double mutant. Conversely, in the 35S::EGL3 and 35S::GL3 overexpression lines, the expression of the CPC::GUS reporter is observed throughout the epidermis in the former case, while in the latter case, there is considerably lower expression along the entire epidermis [16].

Lastly, it has been reported that multiple network components participate in the regulation of GL3 and EGL3 activity. Specifically, when examining the activity of the EGL3::GUS reporter in *wer-1* and *ttg1* mutants, it is observed that there is ectopic expression of this reporter in N-position cells. This observation suggests that WER and TTG1 act as negative regulators of EGL3 expression [17]. On the other hand, in the *cpc-1* and *try-1* mutants an alteration in the expression pattern of EGL3 is not observed, while in the double mutant *cpc-1 try-1* the expression is almost completely reduced. Otherwise, overexpression of CPC and TRY in the 35S:CPC and 35S:TRY mutant lines results in an ectopic expression of EGL3 in N-position cells [17]. Additionally, in the *gl2-1* and *rhd6* mutants, the expression pattern of the EGL3::GUS reporter is indistinguishable from that of the wild type [17]. Finally, in the *gl3 egl3 EGL3::GUS* double mutant, a strong expression of GUS is observed in the N-position cells. Moreover, when either bHLH gene was overexpressed using the 35S::GL3 and 35S::EGL3 constructs, there was a slight to moderate decrease in EGL3::GUS expression. These findings indicate the presence of autoregulation in EGL3 expression, where the EGL3 protein, in conjunction with GL3, is capable of inhibiting the promoter activity of its own gene [17]. The aforementioned findings also apply to the GL3 gene. In the *wer-1* mutant, *gl3 egl3* double mutant, and 35S::CPC overexpression line, ectopic expression of GL3::GUS is observed. Conversely, in the *cpc try* double mutant, a significantly reduced expression of GL3::GUS is observed. The expression of this reporter is unaffected in the *gl2-1* and *cpc* mutants. These observations suggest that the regulation of GL3 closely resembles that of EGL3 [17]. Together this evidence suggests that the activity of GL3 and EGL3 is regulated negatively by the elements that form the activator complex while CPC and TRY do so in a positive way [17].

The CAPRICE (CPC) gene is responsible for encoding a single R3 MYB repeat that does not possess the usual transcriptional activation domain [134] On the other hand, CPC contains a conserved bHLH-binding domain that enables its association with GL3/EGL3. However, it is unable to bind to promoter regions or initiate transcription [123, 145]. A mutation in the CAPRICE (CPC) gene leads to a decrease in the number of root hairs, suggesting that CPC serves as a positive regulator of root hair development [133, 134]. Furthermore, CPC exhibits the ability to move between cells, specifically from N-position cells to H-position cells [134]. The *cpc-1* mutant exhibits elevated GL2 expression, including ectopic activity in certain H-position cells, resulting in a decrease in the overall number of root hairs. This effect seems to rely on the activity of WER, as there is no increase in GL2 expression observed in the *cpc-1 wer-1* double mutant. Conversely, when CPC is overexpressed in the 35S::CPC and 35S::CPC *cpc-1* lines, nearly all epidermal cells develop into hairs [73, 133, 134]. In the *wer-1*, *gl3-1* mutants, as well as in the *gl3-1 egl3-1* double mutant, a significant reduction in CPC expression is evident, with an accumulation of CPC RNA specifically in N-position cells. Conversely, when both genes are overexpressed in the 35S::GL3 and 35S::EGL3 lines, CPC expression occurs in H-position cells [16, 73]. On the other hand, the *wrky75* mutants exhibit an increase in the number of root hairs in the N position. Similarly, these mutants show an elevation in CPC expression in H-position cells. Conversely, the overexpression of WRKY75 in the 35S::WRKY75 line reduces the transcriptional levels of CPC. The *cpc-2 wrky75-25* double mutant displays a phenotype similar to the *cpc-2* mutant. However, when WRKY75 is overexpressed in the *cpc-2* mutant background, there is a lower number of hairs compared to the *cpc-2* mutant alone. This suggests that WRKY75 regulates the differentiation of atrichoblasts and trichoblasts through mechanisms beyond solely suppressing CPC expression. Additionally, the analysis of the CPC gene promoter region reveals that WRKY75 binds to it. Overall, these results demonstrate that the activity of CPC is negatively regulated by WRKY75 [96].

GLABRA 2 (GL2) is preferentially expressed in N-position cells due to differential accumulation of the MBW complex between H and N positions. GL2 encodes a homeodomain-leucine-zipper transcription factor, and because it is produced preferentially in N-position cells and because mutant deficiencies in GL2 promote ectopic root hair growth in N-cells, GL2 is assumed to be a negative regulator of root hair development [32, 81, 95]. A number of bHLH genes, including ROOT HAIR DEFECTIVE 6 (RHD6), RHD6-LIKE 1 (RSL1), RSL2, Lj-RSL1-LIKE 1 (LRL1), and LRL2, have their expression suppressed by the GL2 protein when it binds directly to their promoter regions [77]. Thus these transcriptional factors are responsible for the initiation and elongation of root hairs, such that GL2 expression at the N-position inhibits hair formation resulting in the determination of non-hair cells [82, 84].

Because GL2 is essential for the development of non-hair cells, mutations that affect GL2 expression lead to aberrant root epidermal cell patterns. In the *gl2-1* mutant the roots show a percentage close to 100% of hairs giving rise to a very hairy phenotype [81], in the case of the *wer-1* and *ttg1* mutants, which are unable to form the MBW complex and therefore the GL2 activity is down, very hairy phenotypes are observed due to the increase of hairs in the N-position [43, 72]. Finally, in the case of the *gl3-1* and *egl3-1 m*utants, both exhibit decreased GL2 expression and hairy phenotypes, with the *gl3-1* phenotype being the most drastic, while the *gl3-1 egl3-1* double mutant shows a complete absence of GL2 expression and an extremely hairy phenotype [16]. On the other hand, in the *cpc-1* mutant in which the inhibition of the MBW complex is almost null in the H-position, the ectopic expression of GL2 [133] is observed, showing a marked decrease of hair cells in the H-position in a percentage close to 70%, resulting in a hairless phenotype [133]. On the other hand, in the double mutants *cpc-1 try-82* or *cpc-1 etc1-1* these phenotypes are enhanced and we can observe completely hairless phenotypes [61, 116]. Consequently, this shows that GL2 is an important regulator of atrichoblast identity and any alteration in its expression will modify its proportion.

Another new component is the transcription factor ZINC FINGER PROTEIN 5 (ZFP5) is also involved in the regulation of CPC activity. In the *zfp5-4* mutant and in the ZFP5 RNAi transgenic line, a reduced number of root hairs was observed. On the one hand, a decrease to 75% and 77.6% respectively of the hairs in the H position is observed, which shows that in both mutant lines, there is a significant suppression of the hairs with respect to the wild-type phenotype [2].On the other hand, in the *zfp5-4* mutant, a decrease in CPC and EGL3 levels is observed, suggesting that ZFP5 acts upstream of CPC and EGL3. Overexpression of ZFP5 in the *zfp5-4* mutant is not able to recover the phenotype to that observed in the wild-type phenotype whereas overexpression of CPC in the *zfp5-4* mutant is able to do so. In addition to the above results, the *cpc zfp5-4* double mutant exhibits a similar phenotype to the cpc mutant, while the expression patterns of the CPC::GUS reporter in the *zfp5-4* mutant show that CPC expression is not only restricted to cells in the H-position but in adjacent cells. Finally, by chromatin Immunoprecipitation (ChIP) assay on the 35S:ZFP5:GFP line, it was shown that two regions in the CPC promoter were enriched with GFP antibody, suggesting that ZFP5 directly targets CPC to control root hair initiation. Taken together, these results suggest that ZFP5 acts upstream of CPC [2].

It has been suggested that the TRIPTYCHOME (TRY) and ENHANCER OF CAPRICE AND TRIPTYCHOME1 (ETC1) genes act in a partially redundant manner to CPC in root hair determination [61, 106]. While the CPC, TRY, and ETC1 genes play a role in inhibiting atrichoblast formation, their predominant expression is observed in N-position cells [61, 106, 134], however, diffusion has only been demonstrated for the CPC protein but not for the TRY and ETC1 proteins in the root epidermis [64]. ENHANCER OF CAPRICE AND TRIPTYCHOME1 (ETC1) gene encodes a protein that shares 67% similarity and 54% identity with the CPC protein, which performs a similar role in the differentiation of epidermal cells [125]. In the *etc1-1* mutant a slight decrease in the frequency of root hairs was observed, however, such a decrease is statistically not significant. Conversely, in the 35S::ETC1 line, an abundance of root hairs is observed as a result of the transformation of non-hair cells into hair cells [61]. On the other hand, the etc1-1 try-82 double mutant exhibited a normal quantity and arrangement of both root hair and non-hair cells. However, the double mutant *etc1-1 cpc1* has a significant reduction in root hair production as compared to the *cpc-1*. Finally, the *etc1-1 try-82 cpc-1* triple mutant has a phenotype similar to that of the *try-82 cpc-1* mutant, with the absence of hairs in most of the root being more abundant in the region of the root and hypocotyl junction which is normally a dense zone of hairs. This suggests that ETC1 potentiates the effects of TRY and CPC [61]. It has been suggested that, like CPC, ETC1 has the ability to move from hairless cells to hair cells [108, 133, 136], However, exclusively hairless cells of ETC1:ETC1:GFP transgenic plants showed GFP fluorescence, indicating that the ETC1:GFP fusion protein does not undergo movement within root epidermal cells [125]. In addition to the above result, a strong fluorescence signal is observed in the hairless cells in the CPC:ETC1:GFP transgenic plants, while it is barely detectable in the hair cells; thus both results suggest that ETC1 is unable to mobilize as CPC does [125].

The TRIPTYCHON (TRY) gene encodes a transcription factor that belongs to the MYB family, specifically the single-repeat MYB-related subgroup and whose expression takes place in the cells of N-position [106, 116]. This factor, however, does not possess an activation domain and shows a notable sequence similarity compared to CPC [106]. In *try* mutants, root hair formation is indistinguishable from wild-type phenotype, while a significant increase in the number of root hairs is observed in the 35S::TRY line [106, 116]. Furthermore, when analyzing the expression of the GL2::GUS reporter in the *try* mutant, ectopic expression of the reporter is observed in H-position cells, however, as mentioned, this does not necessarily result in alterations in the pattern since the mutant is phenotypically indistinguishable from the wild-type [116].

As discussed previously, it has been suggested that CPC, TRY and ETC1 act redundantly. For example, the analysis of the *cpc* and *cpc try* mutants shows interesting results: in the first case the mutant shows a considerable decrease of root hairs while in the double mutant, no hairs are observed in the whole root [106]. A detailed analysis with the GL2::GUS reporter in this mutant shows a strong signal throughout the root, which suggests two points, on the one hand, this mutant demonstrates a potentiation of what was observed in the *cpc GL2::GUS* and *try GL2::GUS* lines, and in addition, indicates that CPC and TRY act redundantly in the repression of GL2 in N cells as well as the H cells [116]. RNA expression analysis experiments of TRY in the *wer* and *ttg1* mutants show very little accumulation of its transcript, whereas the *cpc try* double mutant shows a higher accumulation than that observed in the wild-type and the *cpc* and *try* mutants. Thus it can be inferred that TRY is positively regulated by WER and TTG and negatively regulated by CPC and itself [116]. On the other hand, a decrease in TRY transcript levels is observed in the *gl2* mutant; in addition, the expression pattern of the GL2::GUS reporter in the *gl2* mutant shows ectopic expression in the H-position cells similar to that observed in the *try* mutant, so together this evidence suggests that TRY expression is positively regulated by GL2 [116].

The SCRAMBLED (SCM) gene encodes a kinase-receptor-like protein that shows a preference for accumulating on the membrane of cells in the H-position [65,68]. During the early developmental stages in the root epidermis, the SCM-GFP reporter can be observed accumulating in both H-position and N-position cells within the meristematic zone. However, as the development progresses into the late-meristematic and early-elongation zones, the SCM-GFP shows a preference for accumulating in differentiating H cells [66]. The knockout mutants of the SCM gene display a predominantly random accumulation of the WER-GL3/EGL3-TTG1 complex in the root epidermis, as well as root hair patterns that are not dependent on their position [68]. Therefore *scm* mutants alter the distribution of hair and non-hair cells (Kwak et al, 2005). According to previous studies [65], the *cm-2* mutant shows an increase in WER transcription while overexpressing SCM in the 35S::SCM and WER::SCM mutants leads to a notable decrease in WER expression. These findings suggest that the positional signal of SCM involves suppressing WER expression in cells located at the H position. Thus, SCM regulates positional information by reducing the expression of WER in H-position cells [65, 68]. It has been shown that SCM activity is negatively regulated by WER and GL3, and positively regulated by TRY: The expression of SCM::GUS is elevated in the epidermis of *wer-1* and *gl3-1 egl3-1* roots, but decreased in the *cpc-3 try-82* root epidermis. Additionally, the *try-82* single mutant shows a decrease in SCM::GUS expression and presents an abnormal epidermal cell-type pattern [66]. In the *cpc-3 try-82 gl3-1 egl3-1* and *cpc-3 try-82 wer-1* multiple mutants carrying the SCM::GUS reporter, the expression of SCM is notably reduced, similar to the level observed in the *cpc-3 try-82* lines. These findings strongly suggest that the WER-GL3/EGL3-TTG1 complex suppresses SCM expression in N cells, while TRY independently promotes SCM expression in H cells, regardless of the presence of the WER-GL3/EGL3-TTG1 complex [66]. Initially, there was a suggestion that CPC might function as a positive regulator of SCM expression due to its similarities to TRY. However, recent evidence contradicts this segguestion. In the root epidermis of *try-82* mutant plants, the expression of SCM::GUS was significantly reduced. On the other hand, no detectable changes in SCM::GUS expression were observed in *cpc-1* mutants. Interestingly, in the 35S::TRY overexpression line, an increase in SCM::GUS expression was observed, while in the 35S::CPC overexpression line, a decrease in SCM::GUS expression was observed. These findings strongly indicate that TRY, but not CPC, acts as a positive regulator of SCM expression in the root epidermis of Arabidopsis. [67]. While the SCM receptor plays a crucial role in perceiving extracellular positional cues and maintaining the accurate positional patterning of the epidermis [65], it is important to note that the pattern of epidermal organization is established early during embryonic development and does not require SCM. Therefore, it is evident that another mechanism must be involved in the process of determining positional identity in the epidermis,5 JACKDAW (JKD) gene activity being one of these possible mechanisms [51].

JKD encodes for a putative C2H2 zinc finger protein [141]. The *jkd* mutant shows a root epidermis where the hair cell distribution was randomized. A detailed analysis of the *jkd-4* mutant shows that 17% of the cells in the N position produced H cells while about 11% of the cells in position H do not develop as root hairs, suggesting that JKD is necessary for the correct pattern of the epidermal organization [51]. In contrast, when examining the interaction between JKD and the genes involved in the gene regulatory network (GRN) that determines the identity of trichoblasts and atrichoblasts, intriguing information emerges. In the *wer-1* mutant and the *jkd-4 wer-1* double mutant, a hairy phenotype arises due to the formation of hairy cells in N-position cells. Notably, in the double mutant, there is no additive effect between the JKD and WER mutations, indicating that JKD operates through WER to determine this particular phenotype [51]. The same happens when comparing the mutants of *gl2-1* and the double mutant *jkd-4 gl2-1*: the two mutants show very hairy roots as a result of the increase of hairs in the N-position cells and with very similar percentages (67 % and 65.30 %, respectively), suggesting, as in the case of WER, that GL2 acts downstream of JKD in N-position cells [51]. Furthermore, the analysis of the *jkd-4 try cpc-1* triple mutant revealed an identical phenotype to that of the *try cpc* mutant, characterized by a complete absence of H cells. This suggests a genetic epistasis relationship between JKD and the regulators involved in determining the H fate, implying that CPC and TRY function downstream of JKD [51]. Finally, the *scm* and *jkd* mutants share certain phenotypic characteristics: in both cases, the N-position cells can adopt the identity of a hair cell while the H-position cells can adopt the identity of a non-hair cell so that their distribution is not related to their position or to their relationship with the underlying cortex cells. Furthermore, when analyzing the *jkd-4 scm-2* double mutant, an interesting epistatic relationship between *scm-2* and *jkd-4* is observed. In the *jkd-4* mutant, there is a higher percentage of hairs in the H position (90%) compared to the N position (6.8%). Conversely, in the *scm-2* mutant, the percentages are 48.4% and 15.6% for the H and N positions, respectively. However, in the double mutant *jkd-4 scm-2*, the percentages shift to 53.8% for the H position and 17.8% for the N position. These results indicate that SCM acts downstream of JKD in determining the distribution of hairs in different positions [51]. Previous studies have revealed that while JKD transcript and protein are not expressed in the epidermis [141], it is likely that adjacent tissues express JKD, thereby generating the activation signal towards SCM. This is supported by experimental evidence showing that in the *jkd-4* mutant when the JKD gene is introduced under the control of the constitutive cortex-specific pCO2 promoter (pCO2::JKD::GFP), normal hair distribution patterns can be restored. These findings strongly suggest that JKD exerts its influence non-autonomously from the cortex to regulate position-dependent epidermal patterning [51]. The above results suggest that JKD acts in the cortex as a positional information signal for epidermal cells, so a cell in position H located between two cells in the cortex having a larger contact surface is able to receive a larger signal, while cells in position N receive a smaller amount. In this way, there is greater SCM-mediated inhibition of WER in H-position cells and thus they adopt the hair identity, allowing position-dependent determination [51]. Another important new element incorporated into the structure of this network is the WRKY75 gene (WRKY75) [29, 96]. WRKY75 is part of the WRKY family of transcription factors whose role has been described as important in the response to different biotic and abiotic stresses and which can act as transcriptional activators or repressors in various combinations of homodimers or heterodimers [97]. It has been described that WRK75 increases its expression in response to phosphate deprivation in the medium, suggesting that it is an important regulator of root response to alterations in phosphate concentrations [29].

Hormonal signals have a significant role in controlling the genetic machinery that determines root hair differentiation and there is considerable evidence to support this idea [146]. For example, it has been reported that the application of 1-naphthylacetic acid (NAA), an auxin analog, increases the number of cortex cells, which increases the number of hair cells and consequently the density of root hairs due to the existence of more cells in H-position [90], on the other hand, the application of NAA in plants under standard growth conditions produces an increase in the expression of CPC, TRY and ETC1, which generates a decrease in the expression of GL2 due to a decrease in the formation of the MBW complex [74, 90]. On the other hand, the application of N-1-naphthylphthalamic acid (NPA), an inhibitor of polar transport of auxin, results in increased expression of WER, GL3, GL2, and TTG1. These data together suggest that auxins regulate trichoblast identity by increasing the expression of positive regulators of trichoblast identity [74, 90]. In the case of ethylene (ETH), this hormone is involved in both the process of differentiation and determination and in the process of initiation of root hair growth [74] and this happens through the activity of ZFP5 [2]. As was explained, in the *zfp5-4* mutant, a lower root hair density is observed compared to that observed in the WT [2], on the other hand, in the *etr1-1* and *etr1-3* mutants, which are insensitive to ETH, ZFP5 expression is decreased while in the *eto2* mutant, which presents overproduction of ETH, it is increased [2, 53, 74], suggesting that ETH participates in determining the identity of root hairs by regulating the expression of ZFP5 [74]. Furthermore, it has been described that by altering the GRN structure underlying root epidermal differentiation, the effect of exogenous application of ETH is also altered, which can be clearly seen in the *cpc try* mutant because, unlike the *cpc* mutant, exogenous ETH treatment does not recover the density of hairs in the epidermis suggesting that the induction effect of ETH is dependent on the integrity of CPC and TRY [74]. Cytokinin (CK) is another important hormone in the process of determination of *Arabidopsis* root hairs, but the exact mechanism of its functioning is not yet clear [146]. It has been reported for example that its activity is mediated by ZFP5 [2], thus in plants treated with the synthetic CK 6-benzylaminopurine (6-BA) they present an increase in the expression of ZFP5 and a higher density of hairs [2], while in the mutant *zfp5-4* the exogenous treatment with 6-BA is unable to recover the length of the root hairs, suggesting that ZFP5 is necessary for the signaling of CK during the determination and elongation of the hairs of the epidermis of the root [2].

With this information, the network’s topology shown in Figure 2 was established. With this network and the rules defined for each of its interactions previously defined and synthesized in the supplementary information (See Supplementary Information), the dynamic behavior of the network was evaluated. The results obtained are described in detail in the following sections.

### The meta-GRN recovers the spatial organization patterns of different mutants

To verify the relevance of the interactions proposed in the discrete GRN model, a two-dimensional spatial model was established, composed of a 20×20 cell grid that simulates the surface of the epidermis, and *in silico* experiments were carried out on the loss and gain of function of the genes considered in this network model update (See methods for more details). For each simulation the system was initialized with random initial conditions in all elements except for the following cases: TTG1=1, WRKY75=1, and MYB23=0; in the case of hormones a value of 1 is considered as standard growth conditions, such that AUX=1, CK=1, and ET=1.

In order to determine if the meta-GRN reaches a stable state we proceeded to analyze the expression state of the genes GL2, WER, MYB23, GL3/EGL3, TTG1, TTG2, CPC, TRY, ETC1, SCM, ZFP5, WRKY75, as well as the proteins CPC and GL3/EGL3 maintain their stable state for at least the last 100 iterations out of a total of 1000 in 10,000 simulations of the WT phenotype. All genes except GL3/EGL3 maintain steady-state expression in all cells for the last 100 iterations out of 1000 for the 10,000 simulations performed, in the case of GL3/EGL3 they maintained a steady-state for the last 100 iterations only in 9000 out of 10,000 simulations, while in 1000 simulations they reached a steady-state in all cells in the last 100 iterations. Particularly important to note is that the fact that GL2 reaches stability in all cells of the simulated domain implies that the organization patterns of hair and non-hair cells are also stable, suggesting that the network is stable enough to achieve the organization pattern corresponding to the WT phenotype.

In the case of the GL3/EGL3 proteins, these only remained stable after the last 100 iterations out of a total of 1000 in 10,000 simulations, while the CPC protein did not reach a stable state in any of the simulations, which may be a result of the regulation of its expression within the network resulting in greater variability in its concentration in different simulations. Both results suggest that the meta-GRN maintains stable behaviors over time for all the regulators considered even when the concentration of the CPC protein is not stable, and given that the GL2 pattern remains stable and therefore, the identity of the atrichoblasts and trichoblasts determined by their expression level, we can establish that the network is robust and its dynamic behavior is adequate to simulate the expression profiles of the elements considered.

The next step was to characterize the attractors of the system. This network is composed of 13 elements representing transcriptional factors, 1 node representing a transcriptional complex, and 3 nodes representing hormones. Of these regulators, 11 are variables that can take two levels of expression (0 or 1) and 6 variables can take three levels of expression (0, 1 or 2) (See Supplementary Information for more details). Thus the state space is very large and doing an exhaustive search is computationally expensive. To avoid these drawbacks, they first performed about 5,000 simulations starting with a random expression profile for each of the regulators in each of the cells in the simulated cellular domain to retrieve all the resulting attractors, although this does not ensure the recovery of all of the network’s attractors, it is possible to obtain the most common ones or those with the highest basin of attraction. Table 1 shows the recovered attractors in the simulated cell domain of 400 cells and the proportion of each of them for the 10,000 simulations. It can be seen that the configuration with the highest proportion corresponds to the atrichoblast profile while the second configuration corresponds to the atrichoblast profile, the proportion of the other 3 configurations represents approximately 4.3%, of these configurations two attractors have their occurrence in N-position cells (attractors 4 and 5) suggesting that these expression profiles correspond to the gene expression profile of ectopic trichoblasts and in which differences in the CPC expression profile are observed with respect to the H-position trichoblast attractor.

**Table 1.** Attractors recovered in the simulations.

| GRN element | Attractor<br>1 | Attractor<br>2 | Attractor<br>3 | Attractor<br>4 | Attractor<br>5 |
| --- | --- | --- | --- | --- | --- |
| GL2 | 1 | 0 | 0 | 0 | 0 |
| WER | 2 | 0 | 0 | 2 | 2 |
| MYB23 | 1 | 0 | 0 | 0 | 0 |
| GL3/EGL3 (RNA) | 1 | 2 | 2 | 2 | 2 |
| GL3/EGL3 (Protein) | 2 | 2 | 2 | 2 | 2 |
| TTG1 | 1 | 1 | 1 | 1 | 1 |
| TTG2 | 1 | 0 | 0 | 0 | 0 |
| CPC (RNA) | 1 | 0 | 0 | 0 | 0 |
| CPC (Protein) | 2 | 1 | 2 | 1 | 2 |
| TRY | 1 | 0 | 0 | 0 | 0 |
| ETC1 | 1 | 0 | 0 | 0 | 0 |
| SCM | 0 | 1 | 1 | 0 | 0 |
| Proportion | 64.15 % | 31.59 % | 3.41 % | 0.79 % | 0.065 % |
| Probable cell type | A | T | T | T | T |

Consequently, the next step was to analyze the expression profile of attractors 1 and 2 observed in Table 1 and compare them with the expected expression profiles for trichoblasts and atrichoblasts according to the experimental evidence. Table 2 shows on the right side the attractors corresponding to the trichoblast and atrichoblast cell profiles obtained in the simulations while on the left side the expected profiles based on the experimental evidence reported in the literature. As shown, the profiles are similar for the genes considered necessary to define the identity of each cell type, suggesting that the results of the proposed model are consistent with the experimental results for the expression profile of the WT phenotype.

**Table 2.** Reported and simulated expression profiles of GRN elements that characterize trichoblasts and atrichoblasts in the root epidermis.

| Gene or protein | Expected attractors |  |  | Recovered attractors |  |
| --- | --- | --- | --- | --- | --- |
|  | Trichoblast | Atrichoblast | Reference | Trichoblast | Atrichoblast |
| GL2 | 0 | 1 | [32,81,95] | 0 | 1 |
| WER | 0 | 2 | [65,68,72] | 0 | 2 |
| MYB23 | 0 | 1 | [59] | 0 | 1 |
| GL3/EGL3 | 1 | 0 | [17] | 1 | 0 |
| CPC | 0 | 1 | [133] | 0 | 1 |
| TTG1 | 0 | 1 | [18,135]<sup>1</sup> | 0 | 1 |
| TTG2 | 0 | 1 | [58] | 0 | 1 |
| TRY | 0 | 1 | [106] | 1 | 0 |
| ETC1 | 0 | 1 | [61] | 0 | 1 |
| SCM | 1 | 1 | [66,67] | 1 | 0 |
| JKD | 0 | 1 | [51] | 0 | 1 |
| WRK75 | 1 | 0 | [96] | 1 | 0 |

Once the expression profiles obtained were compared with those reported in the literature, the next step was to obtain the organization patterns of the WT phenotype and different loss-of-function mutants reported in the literature for the elements considered in the network. For each of them, we obtained both the spatial pattern and the proportion of trichoblasts and atrichoblasts in each of the cell positions (H-position and N-position). In the Table 3 the values reported in the literature for different mutants and lines of overexpression are shown. It shows the percentage of trichoblasts and atrichoblasts measured in each cell position and the references that support them. On the other hand the Table 2 shows the percentages of trichoblasts and atrichoblasts in each of the cell positions obtained through the simulations of each of the genotypes considered. In the following paragraphs, the results of the spatial patterns and the percentages of each cell type in both positions will be described one by one compared to what is reported in the literature.

**Table 3.**
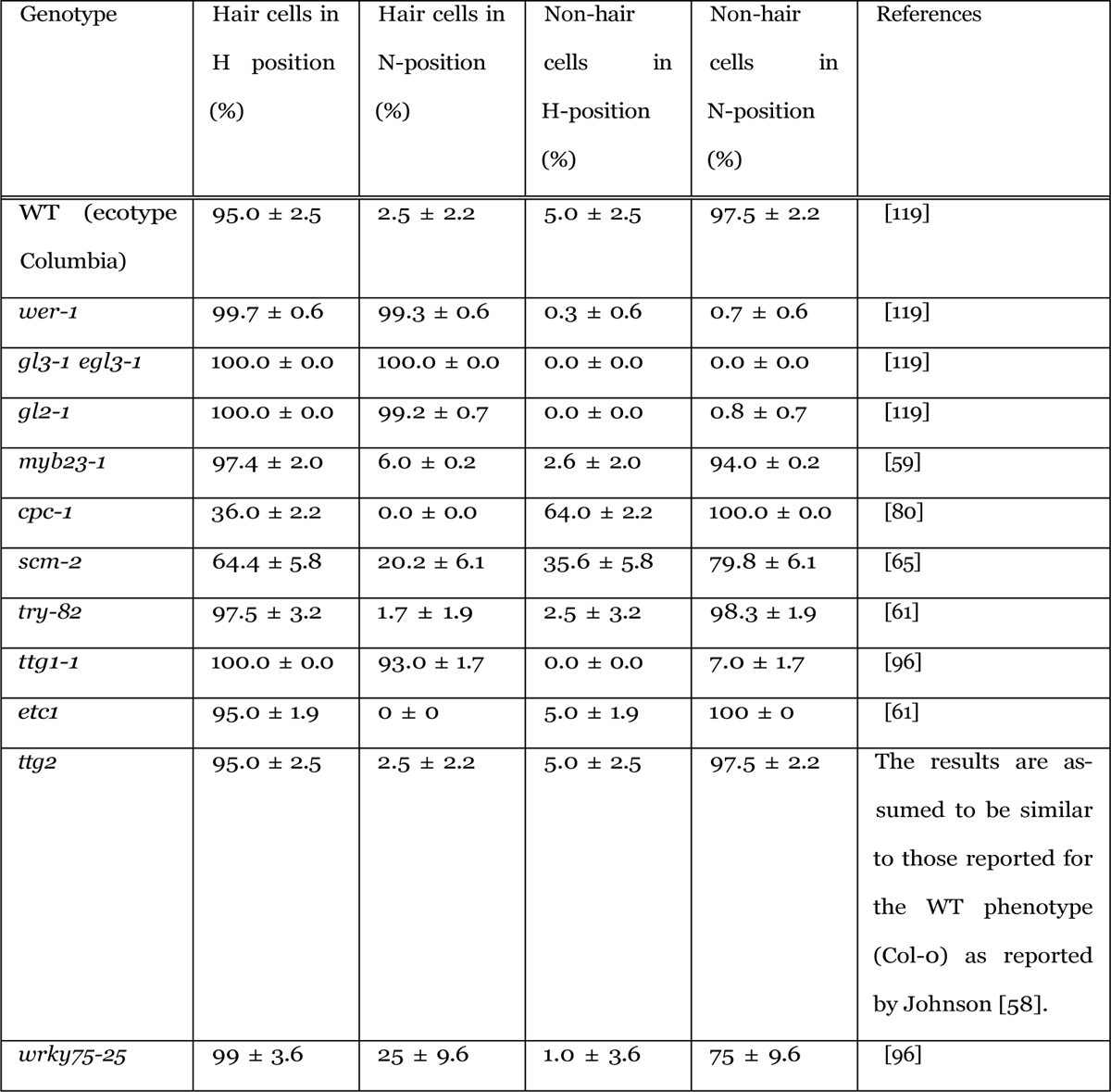
Specification of Cell Types in the Root Epidermis.

In the WT genotype, there are eight root-hair files and approximately 10 to 14 non-hair cell files [37, 43]. This allows expression in an approximate 1:2 ratio between trichoblasts and attrichoblasts. The measurements show that approximately 95% of the cells in the H-position develop as hair cells while a percentage of 2.5% of the cells in the N-position develop as ectopic hairs. On the other hand, 97.5% of the cells in the N-position develop as atricoblasts while a percentage of 5% of the cells in the H-position develop as ectopic atricoblasts. The data from the simulations show similar patterns. On the one hand, there are 2 cell files that adopt an atricoblast expression profile for each row of cells that adopt the trichoblast profile, and on the other hand, the percentages obtained are very close to those reported in the Table 1. These results show that the model qualitatively and quantitatively recovers the spatial organization pattern of the WT genotype.

In the *wer* mutant most of the root epidermal cells differentiate into root hairs through the ectopic production of root hairs. It has been described that more than 90% of cells in the N-position produce root hairs compared to the 5% reported in the wild type [72]. When comparing the results of Table 1 with the results reported in Table 2 we can see that the percentage of hairs in the H-position and the N-position are very similar since while the values reported for the *wer-1* mutant we have 99.7% and 99.3% respectively, the data obtained in the simulations are 100% for both positions. On the other hand, the atrichoblast values both in the simulations and in that reported in the literature are quite similar, a more dramatic phenotype is observed in the simulations since the identity of the atricoblasts is totally reduced. These results suggest that the model is capable of qualitatively and quantitatively recovering the spatial pattern of distribution of trichoblasts and atricoblasts reported for this mutant.

As far as the *gl3 egl3* double mutant is concerned, it produces an extremely hairy root phenotype because of a dramatic reduction in the frequency of the non-hair cell type throughout the root [16]. The data reported for the double mutant *gl3-1 egl3-1* in the upper region of the root show a significant increase in the number of trichoblasts since roots are observed with %98.3 *±* 4.7 hairs in the H-position and %100.0 *±* 0.0 in N-position while the atrichoblasts in N-position disappear. On the other hand, that same mutant in the lower region of the root shows an occurrence of trichoblasts in H-position and N-position of %100 *±* 0 and %96.7 *±* 5.0 respectively, while the atrichoblasts in N-position decrease to % 3.3 *±* 5.5 [16]. On the other hand, when comparing with the results obtained in the simulations, the *gl3 egl3* mutant presents %100.0 *±* 0.0 trichoblasts both in the H-position and in the N-position, while in it presents a total absence of atrichoblasts in both the H-position and the N-position. Although these results are more dramatic, they are similar to those reported for the *gl3-1 egl3-1* mutant in the upper portion of the root, on the other hand, it is observed that the hairy phenotype is the result of an increase in trichoblasts in the N-position due to the absence of atrichoblasts in that position, which corresponds to that reported in the literature. The homozygous *gl2* mutant plants present an excessive root-hair formation phenotype. Root hairs form on epidermal cells in all files and are not limited to epidermal cells located over a radial cortical cell wall. Nearly every epidermal cell in the *gl2* mutant roots produced a root hair [82]. The data reported for the *gl2-1* mutant show that the percentage of trichoblasts in the H-position is %100.0 *±* 0.0 while in the N-position it is %99.2 *±* 0.7, on the other hand the percentage of atrichoblasts in the N-position is of %0.8 *±* 0.7 while in the H-position they are absent [119]. The percentages obtained by simulations of the *gl2* mutant for trichoblasts both in the H-position and in the H-position are %100.0 *±* 0.0, while the percentages of atrichoblasts are %0.0 *±* 0.0 in both positions. These results are consistent with that reported in the literature and show that the phenotype obtained results from an increase in the presence of trichoblasts in the N position. The *myb23* mutant lines appear to have a normal root hair density, but a quantitative analysis of the mutant alleles *myb23-1* and *myb23-3* show a small but statistically significant increase in hair cell production in the N position [59]. As shown in Table 2, if the percentages between the *myb23-1* mutant and the wild-type are compared, a slight increase in the percentage of hairs in the N-position is observed, while the values of hairs and no hairs in both positions are very similar between both genotypes. Therefore, it can be concluded that the model recovers the organization patterns and proportions of atrichoblasts and trichoblasts in both cell positions in a manner consistent with the results reported for the *myb23-1* mutant.

The loss-of-function mutant of CPC shows only a few normal-shaped root hairs and presents a partial decrease in the number of hairs in the H-position and the sporadic occurrence of ectopic trichoblasts in the N-position is not observed [133, 134]. In the *cpc-1* mutant, the number of root hairs in the primary root is about one-fourth of that of the wild type [134].

For the *cpc myb23* double mutant the results are very similar to those reported by the experimental evidence (see Supplementary Information S1-3), it can be observed that when comparing the results of the simulations with those reported for the *cpc-1 myb23-1* mutant, there is a decrease in the number of trichoblasts in the H-position, although the proportion of our simulations is 8% lower than those reported, a marked decrease is observed. In the case of trichoblasts in the N-position, our simulation are 9% higher than reported in the mutant, while the proportion of atrichoblasts in the N-position is 9% lower in our simulations than reported in the literature, finally the proportion of atrichoblasts in the N-position is 8% higher in our simulations than reported for the *cpc-1 myb23-1* mutant.

Consequently, although there is a variation in the proportion of trichoblasts and atrichoblasts in each of the cellular positions, the general pattern is recovered correctly, so it can be assured that this mutant is recovered in a general way with certain restrictions.

The *try* mutant does not show a significant change with respect to the organization pattern neither with respect to the number of trichoblasts and atrichoblasts nor with respect to the number of ectopic trichoblasts in the N-position compared to WT [61]. In the simulations, it is observed that the number of hairs in the H-position is similar to the WT (100% for the WT and 100% to *try*), and the number of ectopic trichoblast in the N-position is very similar (1.2 % for WT and 1.5% for *try*). If we now compare the values of the simulations with respect to those reported for the mutant *try-82* we can see something very similar. Both the trichoblast values in the H-position and N-position are similar between the simulations and the mutant. On the other hand, while the values of atrichoblasts in position-N are similar in both cases, the values of atrichoblasts in position-H are not, since a slight percentage of occurrence of atrichoblasts in position-H is reported for the mutant *try-82* while in the simulations we could not recover atrichoblasts in the H-position. Together the data suggest similar spatial patterns and percentages between the simulations and the mutant lines reported.

For the double mutant *try cpc* our results are similar to those reported in the literature. It can be observed that for the trichoblasts in the N-position and H-position, the percentage is 0% while for the atrichoblasts in both positions, it is 100% (see Section 2 of the Supplementary Information). These results are the same as those previously reported for the *try-82 cpc-1* mutant which shows a complete absence of trichoblasts in the root except in the root-hypocotyl junction region [61]. In this mutant, the proportion of trichoblasts at the N-position and at the H-position is 0% while the proportion of atrichoblasts at the same positions is 100% [51]. Thus, our simulations are equal to the experimental evidence reported for this double mutant.

In the *ttg1* mutant the root produces an excess of root hairs (a hairy root phenotype). The root hairs in this mutant have the same morphology as in the Wild type. Detailed examination of the *ttg1* mutant shows that root hairs form on epidermal cells in all rows and are not limited to epidermal cells located over a radial cortical cell wall. Transverse sections from the mature portion of the root show that the number of epidermal and cortical cells in the *ttg1* mutant roots is indistinguishable from that of the WT. This evidence indicates that the *ttg1* mutation alters the positional control of root-hair cell differentiation, but it does not affect root-hair formation [43, 70]. If we analyze the results reported for the quantification of the number of hairs and non-hairs in the N-position and the H-position in the *ttg1-1* mutant, we can see that the results are very similar with respect to those obtained through the simulations. On the one hand in the H-position the percentage of trichoblasts reported (100.0 %) and simulated (100.0 %) is the same, on the other hand, in the case of atrichoblasts the increase is more dramatic in the simulations (% 100.0) than that reported for the mutant *ttg1-1* (93.0 *±* 1.7), although in both cases the presence of a considerable increase in the number of hairs in the N-position is observed, which is similar to those reported in the description of the phenotype of this mutant [43]. Finally, the number of atrichoblasts in position-H reported (% 0.0) as in the simulations (% 0.0) are comparable with each other, while the atrichoblasts in the N-position are still observed in the mutant *ttg1-1* (% 7.0 *±* 1.7) while in the simulations they disappear completely. These data suggest that the results obtained in the simulations are very similar to those reported in the literature for the mutant *ttg1-1*.

For the double *cpc-1 ttg1-13* mutant, a strongly hairy phenotype is reported in which the proportion of hairy cells at the H-position and at the N-position are 100% and 71.7% while the proportion of non-hairy cells at the H-position and at the N-position are 0% and 28.3%, respectively [80]. On the other hand, the results of our simulations show that although the pattern obtained is that of a hairy phenotype, the percentages recovered are not completely similar to those reported for the mutant (See Supplementary Information). The proportion of hair cells in the H-position is 100%, similar to that reported experimentally, however, the proportion of hair cells in the N-position is 100% while experimental evidence reports 71.7%. This suggests that the excessively hairy phenotype is due to an increase in ectopic hairs and therefore a decrease in the percentage of trichoblasts to 0% at the N-position, however, the phenotype is so drastic that all cells in the N-position differentiate into hairy cells. Thus, although the pattern of organization corresponds to a very hairy phenotype, it is more drastic than suggested by the experimental evidence, so we can conclude that for the *cpc ttg1* mutant our model only partially recovers the phenotype of the *cpc-1 ttg1-13* mutant.

For the *ttg2-1* mutant, it has been described that it does not present alterations in its phenotype with respect to that observed in the WT phenotype [58]. The results of our simulations show that the proportion of trichoblasts and atrichoblasts in each cell position is similar to that reported for the WT mutant both in the simulations and in the phenotypes reported in the literature. Thus, it is possible to conclude (see Table 3 and Table 4) that the model recovers the phenotype of the *ttg2-1* mutant completely.

**Table 4.**
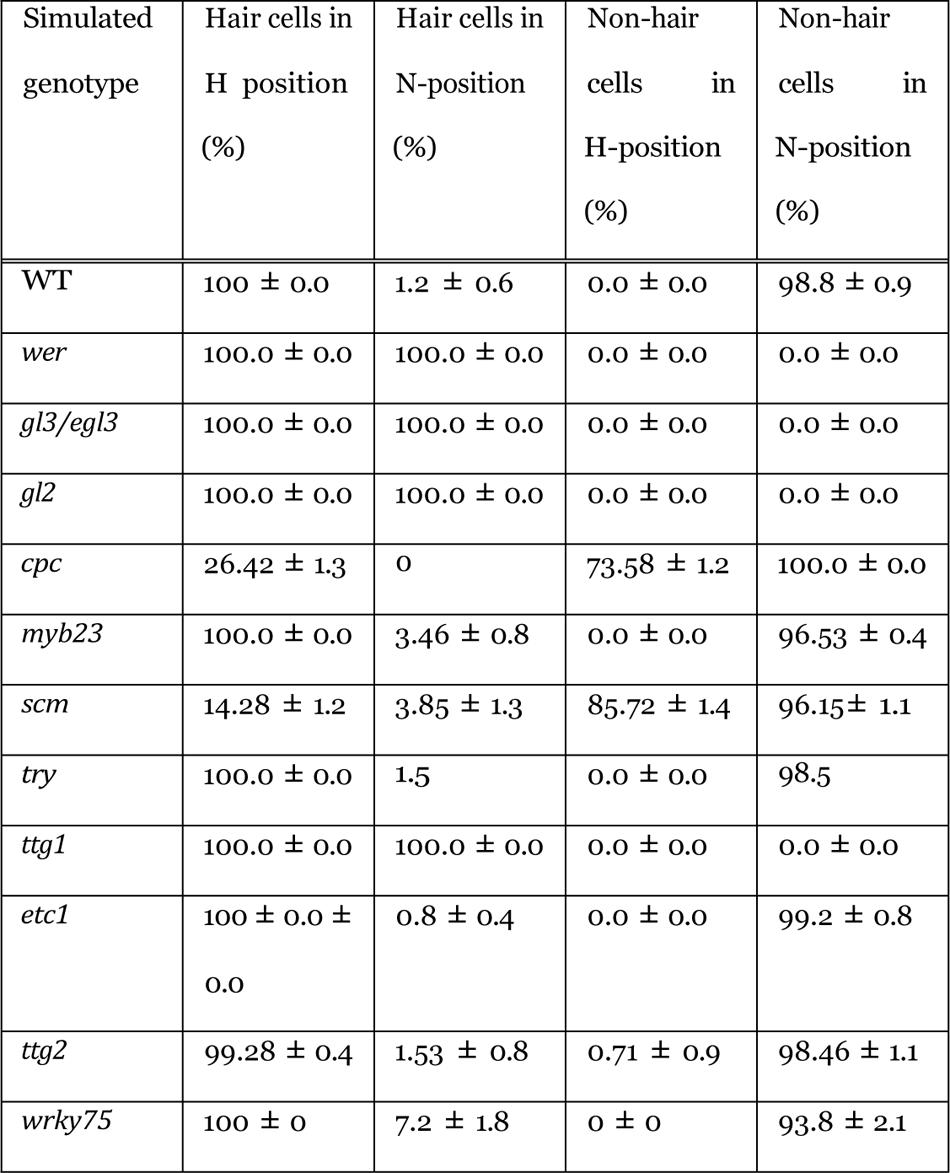
Specification of Cell Types in the Root Epidermis in the simulated domains.

The *etc1* mutant has a slightly reduced frequency of root hairs, but this is not statistically significant and no other abnormalities have been described in the epidermis of the root, which has suggested that ETC1 does not have a central role in epidermal patterning [61]. If we compare the results obtained in the simulation of the *etc1* mutant for the trichoblasts in H-position we have a %100.0 *±*0.0 while the data reported for the *etc1-1* mutant are % 95.0 *±* 1.9, for the trichoblasts in N-position the obtained values are % 0.8 *±*0.6 while those reported for *etc1-1* are % 0.0 *±* 0.0. The atrichoblast values in the N-position and in the H-position are respectively %99.2 *±*0.4 and % 0.0*±*0.0 for the simulations, while those reported for *etc1-1* are % 100.0 *±* 0.0 and %5.0*±*1.9 respectively. Together these data show that the simulations recover values similar to those reported for the *etc1-1* mutant. In the same sense, the double mutant *etc1-1 cpc-1* has been reported to show a significant decrease in the number of root hairs compared to the *cpc-1* mutant [61]. Experimental evidence shows that for this mutant the proportion of hair cells in the H-position and in the N-position are 8.3% and 0% respectively, while the non-hair cells in the H-position and i n t h e N-position are 91.7% and 100% respectively, which shows that in this mutant an increase of non-hair cells in the H-position, *i.e.* ectopic atrichoblasts, is observed. Our results from simulations of the *etc1 cpc* mutant show that the proportion of trichoblasts at the H-position and at the N-positions are 4.3% and 0%, respectively, while the proportion of atrichoblasts at the H-position and at the N-position are 95.7% and 100%. If we compare both results we can notice that in general terms our simulations recover the phenotype reported for this mutant however the proportions are not similar, in particular, our simulations show a higher proportion of atrichoblasts in H-position while the experimental results have a higher proportion of hair cells in H-position (4% difference). Consequently, we can conclude that our model in general recovers the phenotype of the *etc1-1 cpc-1* mutant.

For the etc1 try mutant, our results show only a partial recovery of the pattern with respect to the data reported for the etc1-1 try-82 mutant (see Supplementary Information S1-3). The proportion of H-position trichoblasts in our simulations is 16% higher than that reported for the mutant, while for the number of H-position atrichoblasts our results are 15% lower than that reported for the mutant. Regarding the proportion of trichoblasts in the N-position our results are similar to those reported in the experimental evidence, while for the proportion of trichoblasts in the N-position the difference between the two is 1.56%. These results show that our model partially recovers the behavior of this mutant: the distribution of the atrichoblasts and atrichoblasts in the N-position is recovered, while the proportion of trichoblasts and atrichoblasts in the H-position is not completely recovered.

For the triple mutant *etc1-1 try-82 cpc-1* the experimental evidence reports a phenotype with a total absence of hairs. The percentages reported for this mutant of hair cells in the H-position and in the N-position are 0% in both cases so that the root is completely covered with non-hair cells [61] (See Section 3 in Supplementary data). With respect to the results of our model, our simulations of the *etc1 try cpc* mutant recovers a phenotype completely covered by atrichoblasts, and the proportion of hair cells in H-position and in the N-position are 0% while the proportion of atrichoblasts in both positions are 100%. With this evidence, we can conclude that the model fully recovers the phenotype reported in the literature for this mutant.

In *scm* mutant plants, the arrangement of hair and non-hair cell types does not closely correspond to their typical positions [65, 68]. In the *scm-1* mutant, 66% of cells in the H-position are root-hair cells and 79% of cells in the N-position are non–hair cells. On the other hand, in the *scm-2* mutant, 63.1 *±*8.0 % of the cells in the H-position develop astrichoblasts and 36.9 *±* 8.0 % develop as atrichoblasts, while in the N-position cells, 22.2 *±* 7.3 % develop as trichoblasts and 77.8 *±* 7.3 % as atrichoblasts [68]. Other studies report that in the *scm-2* mutant, in H-position cells 64.4 *±* 5.8 % develop as trichoblasts while 35.6 *±* 5.8 % develop as atrichoblasts, on the other hand in N-position cells 20.2 *±* 6.6% develop as trichoblasts while 79.8 *±* 6.1 % develop as atrichoblasts (See in 2) [65]. From the results obtained by mutant simulations, we can notice two things, on the one hand, the patterns obtained do not show a clear position-dependent organization, so a disorganized pattern of trichoblasts and atrichoblasts is observed, which is consistent with the results reported qualitatively by different studies. Secondly, we note that the number of trichoblasts and trichoblasts in the H-position and in the N-position is lower in our simulations than reported in the literature, while the number of atrichoblasts is higher (see Table 1 and Table 2). Consequently, for this mutant, the model is able to recover qualitatively the disorganized and non-position-dependent pattern of the distribution of trichoblasts and atrichoblasts in the root epidermis but not the total number of trichoblasts and atrichoblasts in each position, so we can say that the behavior of this mutant can only be partially recovered.

For the *scm cpc* and *scm wer* mutants, our results suggest that both are qualitatively and quantitatively recovered by the model when compared to the literature. In the case of the *scm-2 cpc-1* mutant, roots are reported to exhibit a significant reduction in the number of root hairs compared to the *scm-2* mutant, thus in such roots, the expression of the GL2::GUS reporter is markedly expanded and GUS activity is observed in almost all epidermal cells [65]. When comparing the number of trichoblasts and atrichoblasts in the N-position and in the H position, we can observe very similar numbers between those reported in the literature and those obtained in our simulations: although the number of trichoblasts in the H-position is lower than that reported for the *scm-2 cpc-1* mutant, the difference is 1.9%, while the case in the N-position is 0.7%, in the case of H-position and N-position atrichoblasts, differences of 2.2% and 0.7% are observed with respect to the experimental evidence (see Supplementary Information). Consequently, although the number of trichoblasts and atrichoblasts in both positions are different, these numbers are very close to each other and it is possible to establish that both results are comparable.

In the *scm-2 wer-1* mutant, the experimental evidence shows that the roots have an increased frequency of root hairs, a phenotype very similar to that observed in the *wer-1* mutant [65]. If we look at the results of the mutant simulations, the pattern of organization, and the experimental information reported we can notice two things: on the one hand in both cases, an increase in the number of trichoblasts in the epidermis is observed, on the other hand, the comparison of the number of trichoblasts and atrichoblasts has differences (See Supplementary Information). In the *scm-2 wer-1* mutant, the number of trichoblasts in the H-position is 92.1% while in the N-position it is 85.2%, while in the simulated mutant 100% of trichoblasts are obtained in both positions, which indicates that in the case of our simulations, the result is so drastic that all cells differentiate into root hairs, while in the *scm-2 wer-1* mutant this increase is notable but not so drastic so it is possible to observe a small percentage of trichoblasts in position-H (7. 9%) and a higher number of them in the N-position (14.8%); in fact, these percentages are lower than those observed for the *wer-1* mutant (see Table 2). Consequently, it is possible to notice that in general terms our proposed model recovers a similar behavior for this mutant with a more drastic phenotype than the one reported.

For the *wrky75* mutant the results of our simulations suggest a behavior qualitatively similar to that reported in the literature and quantitatively very close to that reported in the literature. As can be observed in Table 2 and Table 3, in the *wrky75-25* mutant 99% of trichoblasts are observed at the H-position while in our simulations it is 100%. For hair cells in the N-position 25% is reported while our simulations report 7.2% of trichoblasts in the N-position, finally, the experimental evidence shows 1% of atrichoblasts in the H-position and 75% in the N-position while our results are 0% and 93.8% respectively in the H-position and N-position (see Supplementary Information). The pattern reported for *wrky75* mutants shows ectopic hairs in the N-position, the function of *wrky75* being to suppress the identity of trichoblasts in the N-position and promote their differentiation in the H-position [96], our simulations recover a similar pattern in which an increase in hair density is observed in the simulated domain that is basically due to the increase of trichoblasts in the N-position, that is, due to the increase of ectopic hairs as expected for this mutant, however, and as can be seen in Table 2 and Table 3, the number of ectopic hairs is lower than that reported for the *wrky75-25* mutant or the WRKY75 RNAi line [96], *i.e.,* our model is able to recover the increase in ectopic hairs of this mutant but not in the proportion reported experimentally.

Our results for the *zfp5* mutant can be seen in Section 3 of the Supplementary Information. Experimental data report that the *zfp5-4* mutant and the ZFP5 RNAi line have fewer and shorter root epidermal hairs than those observed in WT [2]. For the *zfp5-4* mutant, the percentage of root hairs in the H-position and in the N-position is reported to be 75% and 3%, respectively, while the number of non-hairy cells in the H-position and in the N-position is 25% and 97%, respectively. The results of our simulations of the zfp5 mutant show 80.7% and 2.3% for trichoblasts in H-position and N-position respectively, while for atrichoblasts in H-position and in N-position the simulations generate 19.3% and 97.7% respectively. If we compare both results we can see that our simulations are generally similar, although the percentage of trichoblasts in the H-position is 5% higher and 0.7% lower in the H-position, while the number of atrichoblasts in the H-position is 6% lower and 0.7% lower in the N-position, such differences are not so drastic and in general, the recovered pattern shows a decrease of trichoblasts in the H-position leading to a decrease in the whole simulated epidermis. Consequently, we can say that our models are able to recover the patterns reported for this mutant.

The *jkd-4* mutant shows a random pattern in the distribution of hair cells since a 16% increase is observed with respect to the WT phenotype of hair cells in N-position, in addition, 11% of the cells in H-position develop as atrichoblasts, 9% more than that observed in WT, which shows an alteration of the positional determination pattern of trichoblasts and atrichoblasts. If we compare the results of this mutant with the results of the simulations of the *jkd* mutant, we can observe the alteration of the positional determination pattern as expected, however, there are notable differences in the proportions of the two cell types: on the one hand the number of trichoblasts in H-position is 72% lower in the simulations than in the mutant, while the number of atrichoblasts in N-position is 72% higher, on the other hand, the number of trichoblasts in N-position is 3% lower in the simulations than reported for *jkd-4.* Finally, the number of N-position atrichoblasts is very similar to that reported, with a 3.2% higher proportion in the simulations. With this information we can conclude that the mutant partially recovers for the general pattern, the number of trichoblasts and atrichoblasts in the N-position but not for those expected in the H-position.

We must point out that for the *jkd-4 cpc-1* double mutant, the results of the simulations fail to recover the results reported in the literature [51]. This mutant shows a slightly hairy phenotype than that observed for the *cpc* mutant with a marked decrease in the number of trichoblasts in the H-position but less than that observed in the *cpc* mutant and a decrease in the number of atrichoblasts in the N-position, indicating an alteration of the trichoblast/atrichoblasts cell row pattern observed in the WT phenotype. The results of our simulations of the *jkd cpc* mutant (see Supplementary Information S1-3) show a complete absence of trichoblasts in the simulated domain, which is not in agreement with the experimental evidence for the *jkd-4 cpc-1* mutant, making this model unable to reproduce the pattern of organization characteristic of this mutant.

The results of the simulations of the *jkd scm* double mutant show a loss of the typical WT phenotype pattern, a marked decrease in the number of trichoblasts in the H-position, an increase in the number of atrichoblasts in the H-position, and a slight decrease in the number of atrichoblasts in the N-position. If these proportions are compared with the results reported for the *jkd-4 scm-2* mutant, a less drastic phenotype than the simulated one is observed, the loss of trichoblasts is less, so that a percentage of 44% more trichoblasts in H-position is observed, but it is still less than that observed in the WT phenotype, there is a difference of 15. 8% more trichoblasts in N-position, so the percentage of atrichoblasts in the N-position and in the H-position are lower than recovered by the model (see Supplementary Information S1-3). With these results, we can conclude that the pattern of the *jkd-4 scm-2* mutant is partially recovered for the increase of N-position and H-position atrichoblasts, which shows a loss of the pattern of rows of trichoblasts and atrichoblasts that characterize the WT phenotype, but the results are more drastic in the simulations than reported in the literature.

The pattern simulations of the *jkd wer*, *jkd gl2*, and *jkd try cpc* mutants recover very similar patterns to those reported in the literature (see Section 3 of the Supplementary Information). In the case of the *jkd wer* mutant, the simulations show 100% trichoblasts in the H-position and ectopic expression of trichoblasts, while there are no trichoblasts in the H-position and in the N-position. Experimental evidence from the *jkd-4 wer-1* mutant shows that there were 92.5% and 91.6% trichoblasts in the H-position and in the N-position, which are very similar. However, the proportion of H-position and N-position atrichoblasts is 7.5% and 8.4%, *i.e.* the mutant has a low proportion of atrichoblasts in the epidermis, which suggests that in general terms our simulations are consistent with the ectopic increase of trichoblasts, although the simulated phenotype is more drastic, generating hair cells in the whole simulated cell domain. In the case of the *jkd gl2* mutant, the percentage is similar to that obtained for the *jkd gl2* mutant with the whole simulated domain covered with trichoblasts, if we compare with the results reported for the *jkd-4 gl2-1* mutant they show an equal proportion of trichoblasts in position-H and atrichoblasts in position-H, on the other hand, a lower percentage of trichoblasts in position-N and atrichoblasts in position-N are observed (65.3% and 34.7% respectively). This suggests a higher percentage of ectopic hairs in the mutant than what was reported in our simulations, although in both cases the proportion is higher than that reported for the WT, so we can conclude that the pattern is partially recovered, although we were able to recover an increase in the number of ectopic hairs, this proportion is more drastic than that reported for the *jkd-4 gl2-1* mutant, which also shows the absence of atrichoblasts in the N-position.

Finally, in the case of the *jkd try cpc* mutant, the results are identical to those reported for the *jkd-4 try cpc-1* mutant in which a total absence of trichoblasts in the epidermis is observed and, therefore, an increase in the number of ectopic atrioblasts (see Tables 3.1 and 3.2 of the Supplementary Information). Consequently, the model is able to fully recover the pattern and proportion of trichoblasts and atrichoblasts of this mutant.

### The importance of multiple feedbacks in the root hair cell fate establishment

In order to verify the relevance of the proposed interactions and evaluate their effect on the spatiotemporal dynamics of the model, we evaluate the effect on the loss of some of these interactions in the patterns of some mutants as well as in the values of trichoblasts and atrichoblasts obtained. The results of these simulations can be seen in Table 7. A positive feedback loop in WER activity has been proposed as a stabilization mechanism of the pattern of the spatial organization of determination of trichoblasts and atrichoblasts [9, 108]. In previous theoretical works, an auto-regulatory feedback loop of WER activity has been suggested as a key mechanism in the characteristic pattern of organization of the root epidermis without experimental evidence supporting this argument so far. [12, 94] As previously discussed, MYB23 activity represents an important mechanism in regulating the expression pattern of WER [59]. To assess the importance of the previously proposed indirect positive feedback of WER, we simulated meta-GRN dynamics without MYB23 regulation. To compensate for the lack of MYB23 regulation of MBW complex activity, we modified the rules such that the total amount of MYB protein was not altered by the absence of MYB23.

The results of these simulations show some interesting aspects (see Table 7), on the one hand, the WT pattern obtained has a slight decrease in the number of ectopic hairs, and on the other hand, the *scm* mutant has a more drastic phenotype than the one obtained with the meta-GRN that considers the regulation by MYB23, very similar to the results obtained in our previous work [11], even in some of the simulations it is not possible to observe any trichoblast, which suggests that MYB is important in determining the normal pattern of *scm* and to stabilize the pattern of the WT phenotype.

MYB23 expression has been shown to occur at a later stage of root development than in the case of the MYB23 gene [59], this is recapitulated in our model by allowing MYB23 expression to occur with a time delay relative to MYB23 expression. (See the logic rules in the Supplementary Information S1-2). The results for the WT phenotype show that by removing the time delay the WT phenotype loses all ectopic trichoblasts, in the *ttg2* mutant a loss of trichoblasts at the H-position is observed, while in the *scm* mutant no trichoblasts are observed. The *cpc* mutant shows a slight decrease in trichoblasts while the *wer*, *gl3/egl3*, *try*, *gl2*, *ttg1*,*etc1*, *jkd* and *wrky75* mutants show no alteration of the pattern with respect to the simulations where MYB23 time delay was allowed. These results show that the delay in MYB23 expression is necessary to stabilize the WT pattern and the pattern of the *ttg2*, *scm* and *cpc* mutants, with the *scm* phenotype being the most affected by the loss of this temporal regulation, which shows that including the temporal delay in our model is necessary for the description of all mutants and the stability of the WT phenotype pattern. Finally, two interactions are proposed in the model, on the one hand, the direct activation of GL3 and EGL3 by CPC, an interaction proposed from our previous model [11] and suggested in the experimental evidence [16]. To explore the importance of this direct activation, we simulated a version of the network without direct regulation of GL3/EGL3 by CPC and analyzed the formation of the patterns in the WT genotype and the different mutants of the network regulators. When simulating the WT genotype, a pattern similar to that obtained in the simulations of the model with this interaction was observed, however, the trichoblasts in the N-position, which occur occasionally and in low proportion, disappeared completely. On the other hand, the *etc1* mutant loses its ectopic trichoblasts, and the *scm* mutant is devoid of trichoblasts. The other mutants maintain their pattern. These results indicate that direct regulation of CPC on GL3/EGL3 activity is required for the correct determination of the WT phenotype and is necessary to recover the phenotype pattern of the *scm* mutant.

The second interaction proposed in this model is the direct regulation of the TTG2 gene on the activity of the MBW complex. TTG2 has been shown to have promoter binding sites in the TRY gene, physically interacting with TTG1 so that it can bind to the MBW complex, and it appears that this interaction increases TRY expression in the leaf epidermis of [93]. Since it has been shown that the network structure underlying trichome determination shares regulatory mechanisms with the network that determines trichoblasts and atrichoblasts [7, 11], in this work, we have assumed that this regulation also occurs in the root epidermis via regulation of the MBW complex. To assess the relevance of this interaction, we changed the interaction of TTG2 regulation on the MBW complex, instead, we introduced a direct positive regulation of TTG2 to TRY as proposed in the leaf epidermis and with this modified network we simulated mutants of each of the regulators. The results show that the WT phenotype again loses its trichoblasts in the N-position so the pattern shown in the previous results is lost, the *scm* mutant also loses all trichoblasts so a bald phenotype is observed, the *etc1* mutant loses its ectopic trichoblasts, while the *try* mutant phenotype loses some trichoblasts in H-position and all trichoblasts in N-position. This suggests that the regulation of TTG2 on the formation of the MBW complex as proposed in this model is necessary for the correct determination of the phenotype of these mutants, and very importantly for the case of the *scm* mutant, as well as for the correct determination of the WT phenotype. The result of these simulations can be seen in the supplementary information (See Supplementary Information S1-3).

### The importance of diffusion in establishing the pattern of organization of the epidermis of the root

The role of cell communication in establishing the characteristic organization pattern of trichoblasts and atrichoblasts in *Arabidopsis* root epidermis has been widely discussed [11, 103]. As previously described, the GL3, EGL3, and CPC proteins diffuse between the cells of the epidermis and experimental evidence confirms this [17, 60, 64, 73]. We first analyzed the effect of diffusion in determining the pattern of trichoblasts and atrichoblasts in the WT phenotype. To do so, the network was simulated by making changes to the code. On the one hand, the total loss of the diffusion rules was simulated, that is, the network was simulated without considering the diffusion processes. By simulating the dynamics of the meta-GRN they avoid that GL3/EGL3 and CPC proteins diffuse and the state of the nodes depends only on the positional signal of JKD and SCM, *i.e.* by omitting the cell-to-cell communication process, the pattern obtained does not reflect the characteristic organization of rows of WT trichoblasts and atrichoblasts, so that they differentiate independently of their position in the cellular domain (see section 4.1 of the Supplementary Information). These results confirm the initial hypothesis that the pattern of trichoblast and atrichoblast row organization characteristic of the epidermis in WT requires cell-cell communication through the diffusion of GL3/EGL3 and CPC proteins. On the other hand, since JKD and SCM are acting in each of the cells, it is not possible to recover the WT phenotype, suggesting that the positional information of these two regulators alone is insufficient to determine the distribution pattern of trichoblasts and atrichoblasts in the epidermis.

Next, we evaluated the effect of lateral diffusion of CPC, GL3/ELG3 on pattern determination. In our previous model, it was proposed that the elongated shape of the root epidermal cells would result in diffusion in the lateral direction being more important than diffusion in the apical direction allowing this pattern to be more stable and making it possible to recover a lower proportion of ectopic trichoblasts as observed in the WT phenotype [9, 11]. Therefore, to evaluate this effect, we generated a meta-GRN model in which the diffusing elements can only diffuse apically and not laterally. The simulations showed a decrease in the percentage of ectopic hairs with respect to the WT simulations (See Figure 2 in Section 4 of the Supplementary Information). These results are similar to those obtained by our previous model. If we analyze the percentage of ectopic trichoblasts obtained in these simulations with respect to that reported for the WT phenotype, we can see that it is 0.4%, one-third of that reported in the literature. These results suggest that lateral diffusion is important for the establishment of the WT phenotype in which ectopic trichoblast formation is observed sporadically.

Since the above results suggest that positional information via JKD and SCM is not sufficient to allow the correct formation of the pattern of epidermal trichoblasts and atrichoblasts and that diffusion is necessary for this purpose, we evaluate the role of cellular communication through the mobilization of CPC, GL3/EGL3 proteins and tested the effect of the diffusion parameters of these elements in the meta-GRN dynamics and the effect on the pattern obtained. In our previous work, the effect on the variation of the spatial pattern corresponding to the WT was analyzed by varying the diffusion parameters of CPC and bHLH from 0 to 0.25 with a step size of 0.1, observing a percentage error in the pattern of approximately 0.16%, the which can be interpreted as ectopic hairs, a lower percentage than that reported for the Col-0 phenotype [11]. To explore a similar hypothesis, we analyzed the spatial pattern formation effect of trichoblasts and atrichoblasts corresponding to the WT phenotype by exhaustively varying the diffusion parameters of CPC and GL3/EGL3. First, the variation in the total number of hairs in the WT phenotype was analyzed by varying the diffusion value of CPC while keeping the diffusion value of GL3/EGL3 fixed. In this way, the CPC diffusion value was varied from 0 to 1 with a step value of 0.005, and the GL3/EGL3 diffusion value remained fixed at 0.01. For each parameter combination, 100,000 networks were simulated, and then we obtained the total and ectopic root hairs number mean. As observed in Figure 3, by varying the value of the CPC parameter D (Dcpc), it is observed that the variation in the average of the total number of hairs in the cellular domain defines 3 regions of the cellular organization pattern. In the first region that corresponds to the variation of Dcpc from 0 to 0.2916, the average number of hairs oscillates in an average of 140.8-143.6, which is a range similar to that observed in the simulations of the WT phenotype with the proposed parameters for our previous model and in which the mutant phenotypes are recovered semi quantitatively (See Table 4). The second region corresponds to the range of values from 0.2617 to 0.33, where the total number of hairs falls to 123.5 hairs, however, this region is more restricted with respect to region 1 and a fall of 20 total hairs is observed. The last region includes the range of Dcpc that goes from 0.335 to 1, in this region the average total number of hairs varies between 80.7 to 81.9. In this last region, a fall of 62 hairs is observed with respect to the WT phenotype observed in Region 1 and 42 with respect to what was observed in Region 2.

**Figure 3.**
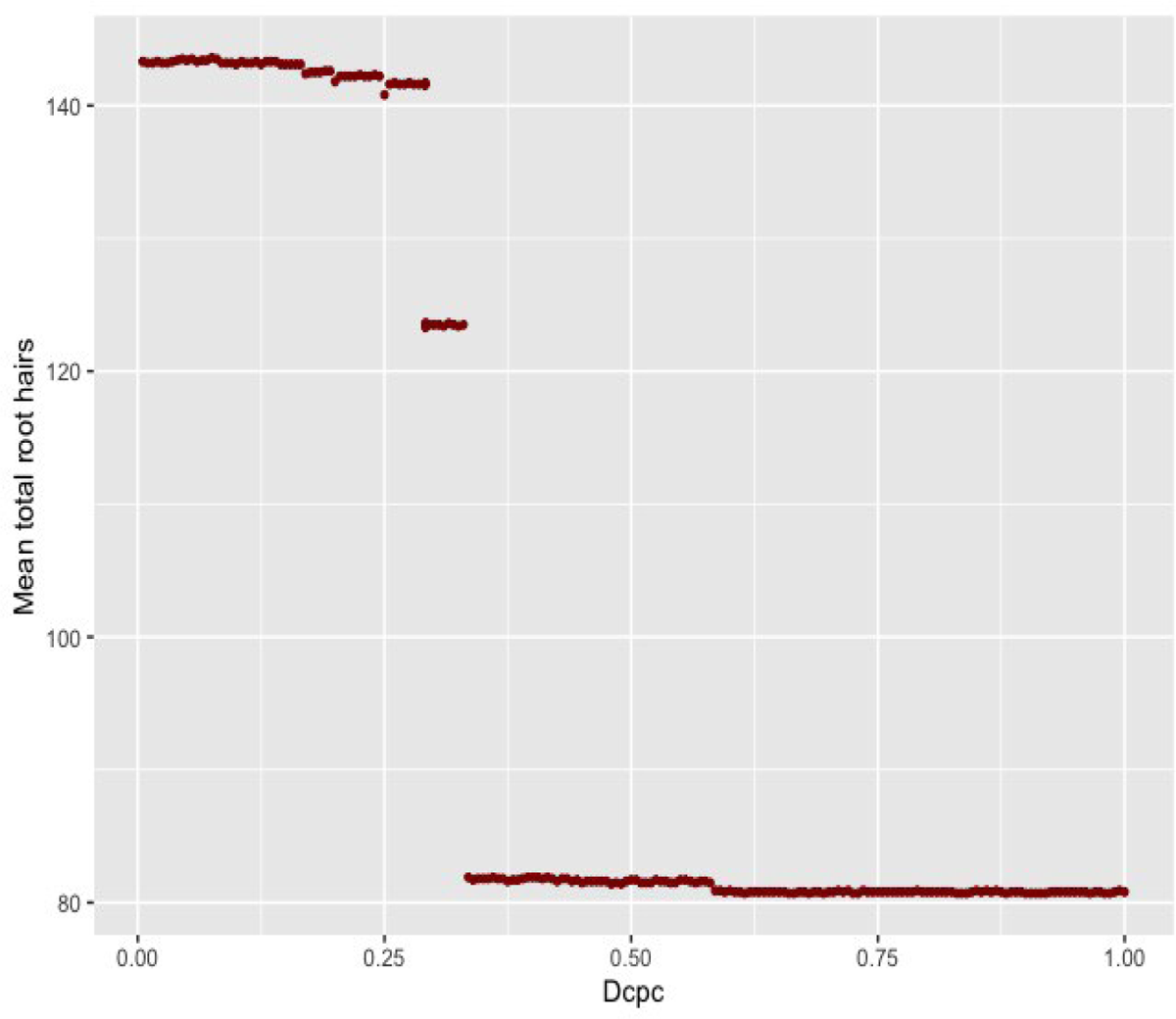
Variation of total root hairs by modifying the diffusion of CPC. The effect of the variation in the Dcpc value on the total number of hairs in the simulated domain is observed. 100,000 nets with Dcpc value were simulated and the average number of hairs was obtained. The average of the total number of hairs in each case is shown.

In order to have a clearer vision of the effect of the variation of the parameter D of CPC in the establishment of the organization pattern of the trichoblasts and atrichoblasts in the context of the proposed meta-GRN, the spatial patterns obtained in the different regions of parameters were previously analyzed. Table 6 shows a sample of the organization patterns of the root hairs obtained for different CPC diffusion values. The first two patterns correspond to the values included in region 1 of Figure 3 (Dcpc=0.05 and Dcpc=0.29), the pattern corresponding to a value of Dcpc=0.30 corresponds to region 2 and the last two patterns (Dcpc= 0.40 and Dcpc=0.80) correspond to region 3. Regarding the patterns of region 1, it is observed that these correspond to what is expected for the WT phenotype reported in our previous work [11] and what is observed is a variation in the number of hairs in position-N, with a tendency to decrease in the hairs in position-N as the value of D increases. Regarding the hairs in the H-position, it is observed that the pattern is not altered and the integrity of the trichoblast rows is maintained throughout the variation of parameter D.

Regarding the pattern corresponding to the value of Dcpc=0.30, which corresponds to region 2, it is observed that the expected organization pattern for the WT phenotype is partially altered. Of the total rows of trichoblasts that are observed in the pattern corresponding to the WT genotype, 28.5% of the rows suffer an alteration in which it is observed that not all the trichoblasts manage to develop, giving rise to patches of atrichoblasts in the rows corresponding to trichoblasts, which is related to the drop in the total number of trichoblasts observed in Figure 3 to an average of 123.5 trichoblasts. Finally, in the patterns corresponding to the values Dcpc=0.40 and Dcpc=0.80, it is observed that the characteristic organization pattern corresponding to the WT genotype has been lost, although it is still possible to observe rows, they have lost their integrity, therefore the number of trichoblasts decreases, however, the number of ectopic trichoblasts increases, being 3 times greater with respect to what was observed in region 1 and almost 6 times what was observed in region two (See Supplementary Information).

The next analysis carried out was the variation of the GL3/EGL3 diffusion parameter while keeping the CPC diffusion value fixed to observe the changes in the number of total and ectopic hairs in the simulated domain. In this way, the GL3/EGL3 diffusion values were varied from 0 to 1 with a step value of 0.005, and the CPC diffusion value remained fixed at 0.05. For each parameter combination, 100,000 networks were simulated, and then we obtained the total and ectopic root hairs number means.

As can be seen in Figure 2 of the Supplementary Information (see section 4.2 of the Supplementary Information), by varying the value of the diffusion parameter D (DbHLH) of GL3/EGL3, it is possible to observe 2 behavioral variations in the pattern of epidermal organization. In the first region corresponding to the DbHLH variation from 0 to 0.495, the average number of total hairs ranges from 143.3 to 145.9, which is similar to that observed in the simulations of the WT phenotype with the parameters proposed in our previous model where an average number of ectopic hairs between 3.3 to 10.7 is observed (See Supplementary Information). The second region corresponds to the range of values from 0.5 to 1, where the average number of total hairs ranges from 200.4 to 201.4, and in this region the increase in the number of total hairs is due to the increase in the number of ectopic hairs in a range from 129.1 to 130.6 on average. In order to have a clearer vision of the effect of the variation of the parameter D of GL3/EGL3 in the establishment of the organization pattern of the trichoblasts and atrichoblasts in the context of the proposed meta-GRN, the spatial patterns obtained in the two regions of parameters.

Table 15 of the Supplementary Information shows a sample of root hair organization patterns obtained for different DbHLH diffusion values. The first three patterns correspond to the values included in the first region of Figure 1 of the supplementary material (DbHLH=0.05, DbHLH=0.25, and DbHLH=0.45), while the last three correspond to the second region (DbHLH=0.5, DbHLH=0.75, and DbHLH=0.99). As seen in the patterns of the DbHLH=0.05 and DbHLH=0.25 values, it can be observed that the distribution of trichoblasts and atrichoblasts is similar to those observed in WT, with well-defined rows of trichoblasts and atrichoblasts and the occurrence of ectopic trichoblasts in a proportion close to 1% and are similar to those reported in our previous work [11]. On the other hand, in the pattern of the DbHLH=0.45 value, the total number of ectopic hairs is on average 3 times larger than that observed for the WT simulations, although the total number of trichoblasts is similar to that observed in WT. When observing the obtained pattern, it can be noted that in spite of conserving part of the WT phenotype pattern, alterations of the same begin to be seen, observing both trichoblasts and atrichoblasts in the N-position and in the H-position, which shows that the increase of the value of the diffusion parameter in the range of 0 to 0.5 increases the number of ectopic hairs but the partial alteration of the normal pattern without altering the total number of trichoblasts. Finally, as can be seen in the last three simulations of Table 15 of the Supplementary Information, having DbHLH values equal to or greater than 0.5, a higher number of total hairs is recovered in the simulated patterns but the recovered pattern in which trichoblasts and atrichoblasts are observed regardless of the position which also leads to an increase of ectopic hairs. This shows that the increase in GL3/EGL3 diffusion values generates an increase in the total number of hairs and this increase is due to the increase in the number of ectopic hairs, however, the pattern is lost and the cells differentiate into trichoblasts and atrichoblasts regardless of the position. This is an inverse behavior to that observed when varying the Dcpc values in which the increase generates a lower number of total hairs due to the loss of the number of hairs in the N-position.

To study the effect of the variation of both diffusion parameters in determining the number of trichoblasts and atrichoblasts as well as their pattern of spatial organization, the joint effect of the variation of Dcpc and Dgl3/egl3 in the determination of the number of hairs was evaluated. Total and ectopic root hairs for combinations of Dcpc and Dgl3/egl3 values from 0 to 1 with a step size of 0.05 were evaluated. For each combination of parameters, 100,000 networks were simulated and the average of total hairs and ectopic hairs was calculated. Figure 4 shows the space formed by the combinations of the Dcpc and Dgl3/egl3 parameters. The color scale shows for each combination of parameters the logarithm of the average of the total number of hairs for 100,000 simulations. As can be seen, the space can be divided into 5 regions with respect to the total number of hairs obtained and their spatial pattern. Region one corresponds to the space contained between the values of Dcpc=0 and Dgl3/egl3=0 to Dgl3/egl3=0.45. As can be seen in patterns A and B within this region, two things are observed: on the one hand, rows of trichoblasts are observed throughout the domain; however, these are incomplete and trichoblasts interspersed with atrichoblasts can be observed. On the other hand, as can be seen in the data shown in Supplementary Information in conjunction with the patterns in Figure 4, the number of ectopic trichoblasts is very low, becoming larger as the Dgl3/egl3 value approaches 0.45. That is, in this region of the parameter space the dynamics of the network are insufficient to determine the totality of the rows of trichoblasts as the ectopic trichoblasts expected for the WT genotype. The second region is found between the values of Dcpc from 0.05 to 0.25 and Dgl3/egl3 from 0 to 0.45. As can be seen in Supplementary Information and pattern C in Figure 4, in this region it is observed that the rows of trichoblasts show their complete identity and the number of ectopic trichoblasts maintains a number similar to that expected for the WT genotype and as the value of Dgl3/egl3 approaches 0.45, an increase in the number of ectopic hairs begins, while as the value of Dcpc approaches 0.25, the number of ectopic hairs decreases. Consequently, this region corresponds to patterns similar to those expected for WT genotypes where complete trichoblast rows are observed interspersed by atrichoblast rows and sporadic trichoblasts in the atrichoblast position. The third region is found between the values of Dcpc=0.3 and Dgl3/egl3=0 to Dgl3/egl3=0.45, which includes pattern D in Figure 4. As can be seen, figure D shows a partially altered pattern with some rows of complete trichoblasts interspersed with rows of atrichoblasts, however, other rows of trichoblasts are incomplete and are interrupted by atrichoblast cells, specifically in the lateral regions of the domain. Consequently, the number of total trichoblasts is less than that observed for the WT pattern, while an average decrease in the number of ectopic hairs is also observed. The fourth region that contains the pattern E is located between the values of Dcpc=0.35 to Dcpc=1 and Dgl3/egl3=0 and Dgl3/egl3=0.4. As can be seen in Figure 4 and the Supplementary Information, a decrease in the number of trichoblasts in the hair position is observed while the number of ectopic hairs on average increases, and as the value of gl3/egl3 increases, so does the value of gl3/egl3. of ectopic trichoblasts. The pattern also shows that it is possible to distinguish separate trichoblast rows from atrichoblast rows even when they are not complete. The fifth region is located between the values of Dcpc = 0.35 to Dcpc = 1 and Dgl3/egl3 = 0.45. As observed in the F pattern, the number of total hairs is greater, this, as observed, is mainly due to the increase in the number of ectopic trichoblasts. On the other hand, it is not possible to locate isolated rows of atrichoblasts since the expression of atrichoblasts is observed in almost the entire domain, showing a pattern similar to a checkerboard with some patches of atrichoblasts. The sixth region is located in the values of Dcpc=0.15 and Dgl3/egl3=0.5 to Dgl3/egl3=0.55. As shown in pattern G, in this small region perfectly defined trichoblast rows are observed separated by atrichoblast rows, however, the number of total trichoblasts is higher due to the increase in the number of ectopic trichoblasts, which is on average ten times more larger than that observed in the WT phenotype characteristic of the second region (see Supplementary Information).

**Figure 4.**
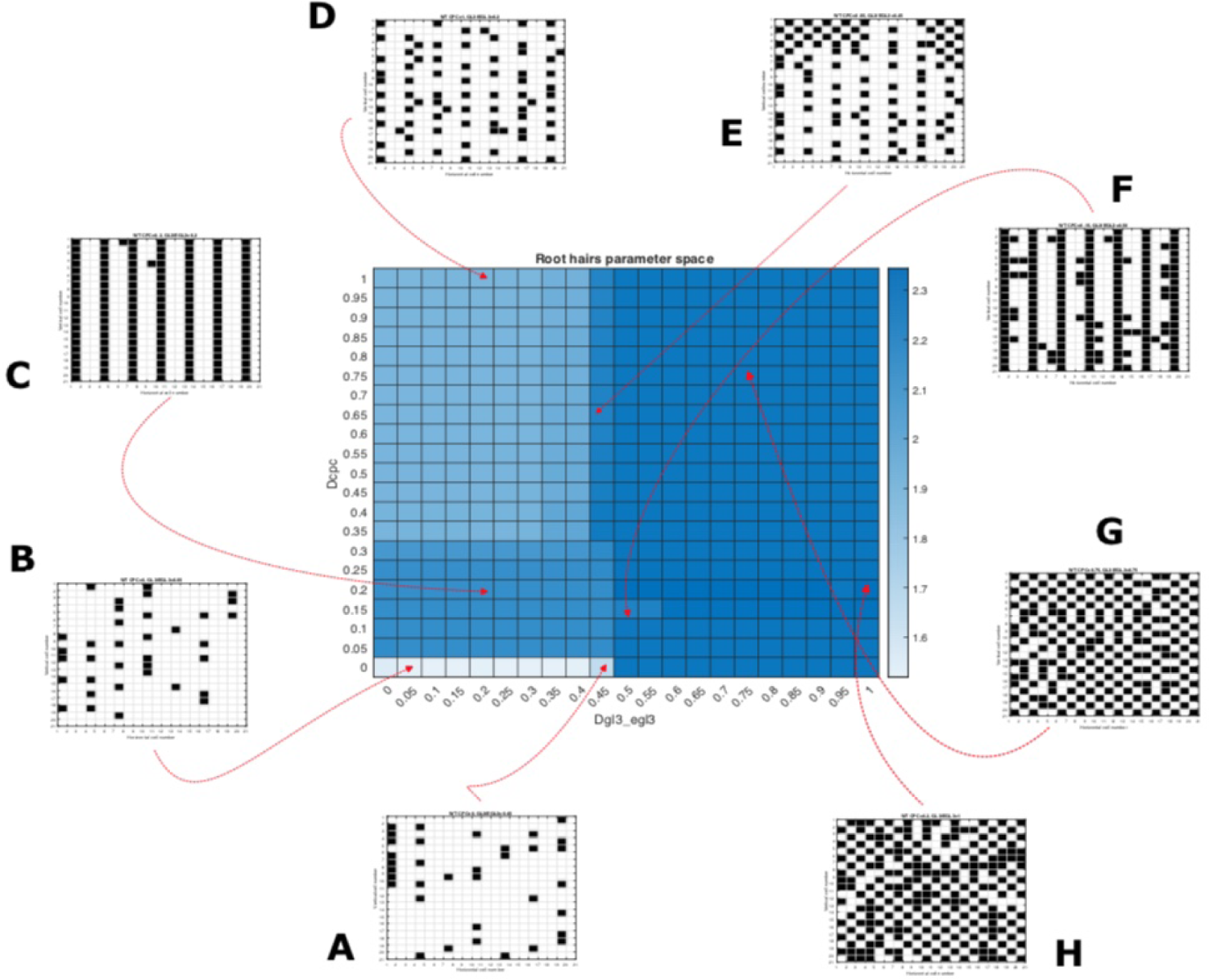
Root hair patterning parameter space. The parameter space consisting of the values of Dcpc and Dgl3/egl3 is shown. The color scale is the logarithm of the total number of trichoblasts in the simulated domain. Eight trichoblast/atrichoblast patterns are shown for different combinations of Dcpc and Dgl3/egl3 in the regions identified in the parameter space. Patterns A and B are found in the first region. Pattern C is found in the second region. The D pattern is found in the third region, the E parameter in the fourth region, the F parameter in the very small region 5 with respect to all the others, and finally the G and H patterns in the sixth region. The patterns located in the second region of the parameter space correspond to those observed for the WT phenotypes of the Arabidopsis thaliana epidermis with variations in the number of ectopic hairs (See Supplementary Information).

The seventh and last region is located between the values of Dcpc=0 to Dcpc=1 and Dgl3/Degl3=0.5 to Dgl3/egl3=1, except within it the seta region previously described. As seen in the H and I patterns, the loss of the characteristic pattern of rows of trichoblasts is observed and in its place patterns resembling chessboards are observed, on the other hand, a considerable increase in the number of ectopic trichoblasts is observed, which is on average 131 against the 3 observed in the pattern corresponding to the WT phenotype.

The above results allow us to conclude three points about the role of diffusion in the formation of the pattern of trichoblasts and atrichoblasts in the root epidermis of *Arabidopsis*. Firstly, results of the simulations in which the diffusion of CPC and GL3/EGL3 proteins was not allowed, it was not possible to recover the WT phenotype, and the pattern structure was completely random, although it was possible to obtain the differentiation of trichoblasts and atrichoblasts these did not have the appropriate position, which shows the importance of diffusion in the dynamics of the GRN and in the establishment of the trichoblast and atrichoblast rows in the epidermis. Secondly, the results of the simulations in which the values of the diffusion parameters of the CPC and GL3/EGL3 proteins were varied show that their alteration is capable of modifying the trichoblast/atrichoblast pattern, in the case of the CPC protein, the increase of the Dcpc parameter value generates a lower number of hairs due to the loss of trichoblasts in H-position and therefore the loss of the integrity of the pattern, on the other hand the variation of the DbHLH parameter shows that its increase generates the total loss of the pattern and differentiates both trichoblasts and atrichoblasts in the H-position and in the N-position which results in an increase in the total number of hairs. Thirdly, by analyzing the joint variation of both parameters it is possible to identify different regions in which the pattern behavior is different, being the C region the one that recovers the phenotype observed in WT, suggesting that the combination of parameters of the WT phenotype is less robust in the diffusion parameter space and thus showing the importance of the proper diffusion of CPC, GL3 and EGL3 proteins for a correct distribution and differentiation of trichoblasts and atrichoblasts in the epidermis.

## Discussion

### Meta-GRN is able to recover *Arabidopsis thaliana* root epidermal organization patterns for different mutant lines

In this study, we introduce a discrete gene regulatory network model that can accurately predict the organizational patterns of trichoblasts and atrichoblasts in the root epidermis for both the wild-type phenotype and a significant number of mutants. Our modeling approach employs as a pint of departure a discrete gene regulatory network that accounts for numerous genetic interactions and simulates the concentrations of epidermal proteins involved in cell-cell dynamics. Through the dynamic behavior of the network, the model can capture the characteristic pattern of rows of trichoblasts and atrichoblasts, enabling cell coupling and the establishment of this organizational pattern. This model is the result of many years of exhaustive functional, genetic, and molecular studies in the *Arabidopspis thaliana* root epidermis.

Although the previous model developed by our research group has been described in detail before [9–13], not all mutant genotypes were examined and only a few were analyzed thoroughly. Nonetheless, our approach to extending that previous model has allowed us to demonstrate its significant explanatory power in recovering the organizational patterns of a broad range of mutants, as described in the literature.

Upon reviewing the results of our simulations for the genotypes considered in this model, we find that they can exhibit four behaviors: those showing changes in the number of trichoblasts/atrichoblasts, those showing changes in the normal position of trichoblasts/atrichoblasts, those showing no changes compared to WT phenotype and finally those showing changes in number and position. Those mutants showing a phenotype in the first category are *wrky75* and *cpc*, those showing a change in position are *scm* and *jkd*, those showing no change are *ttg2*, *try* and *etc1*. Finally those showing both changes in number and position are *wer*, *gl3/egl3*, *gl2*, and *ttg1*. For each of these mutants, the expression patterns in the simulated cell domain and the proportion of trichoblasts and atrichoblasts at the N-position and in the H-position were obtained.

The WT phenotype is completely recovered when compared with the experimental evidence, and in it, columns of trichoblasts separated by two rows of atrichoblasts are observed, and occasionally in the N-position ectopic trichoblasts are observed, the percentage of occurrence of which is less than 2% [37, 43], furthermore, the percentage of trichoblasts and atrichoblasts is similar between simulation results and reported experimental evidence. (Tabla 1 and Table 3 with a variation of 5% for the proportion of atrichoblasts in the H-position previously reported [119]. This demonstrates that the regulators integrated into this model are sufficient to recover the pattern of trichoblasts and atrichoblasts seen in the WT phenotype of the *Arabidopsis* root epidermis in a robust form.

One of the important regulators analyzed in the proposed meta-GRN is WER whose role is central to the atrichoblast identity regulation [72]. As can be seen in Table 5, a pattern similar to that reported in the literature is observed [72, 119]. A detailed review of the values obtained for the *wer-1* mutant shows parameters very similar to those reported by our simulations (see Table 2 and Table 3). However, recently other WER mutants have been described and their alterations in the characteristic root organization pattern do not correspond to the *wer-1* phenotype. For example, the *wer-4* mutant has been identified whose phenotype is less drastic than that observed for *wer-1*. This mutant presents 13% atrichoblasts in the H-position and 28% trichoblasts in the N-position, which contrasts with the strong hairy-root phenotype due to the loss of non hair cells of the other WER mutants. The WER protein of *wer-4* mutant shows a single-residue substitution at position 105 that causes an alteration of the transcriptional patterns and consequently an alteration in the GRN dynamics that prevents the correct determination of trichoblasts and atrichoblasts. Particularly this single-residue substitution alters the WER protein affinity for the GL2 and CPC promoter-binding sites and therefore this alters the cell-type-specific accumulation of the WER-GL3/EGL3-TTG1 complex and consequently alters the accumulation patterns of GL2 and therefore the correct determination of trichoblasts and atrichoblasts [137]. In this sense, our model is not capable of recovering the *wer-4* phenotype since, due to the proposed simulation approach, it is not possible to capture the variations in the ability of the WER protein to differentially bind to the CPC and GL2 promoters. A possible solution to extend the capabilities of this model could be to establish different expression levels for WER, each associated with a WER affinity, and thus capture variations in the levels of WER and the MBW complex capable of determining changes in the expression patterns at the H- and at the N-positions. The *wer-4* mutant displays a unique distribution of cell types in the root epidermis. Unlike the conventional position-dependent cell fate specification, this mutant generates both root-hair cells and non-hair cells in both the H-position and in the N-position [137]. A possible mechanism has been proposed to explain the pattern of this mutant: in *wer-4* the low affinity of WER for the promoters of CPC, TRY and ETC1 generates low levels of the inhibitor complex, there was a normal accumulation of the MBW complex in the H-position and an increase in the levels of the CPC proteins in the H-position, the latter causing animal movements of CPC towards the N-position, which together alter the formation of the MBW complex in both positions. Since the effect of SCM is present at the N-position, there may be a relative increase of the MBW complex. Thus this whole process generates an ectopic expression of GL2 so that the determination of the identity of the atrichoblasts or trichoblasts depends on the accumulation of GL2 and not so much on the position of the MBW complex [137].

**Table 5.**
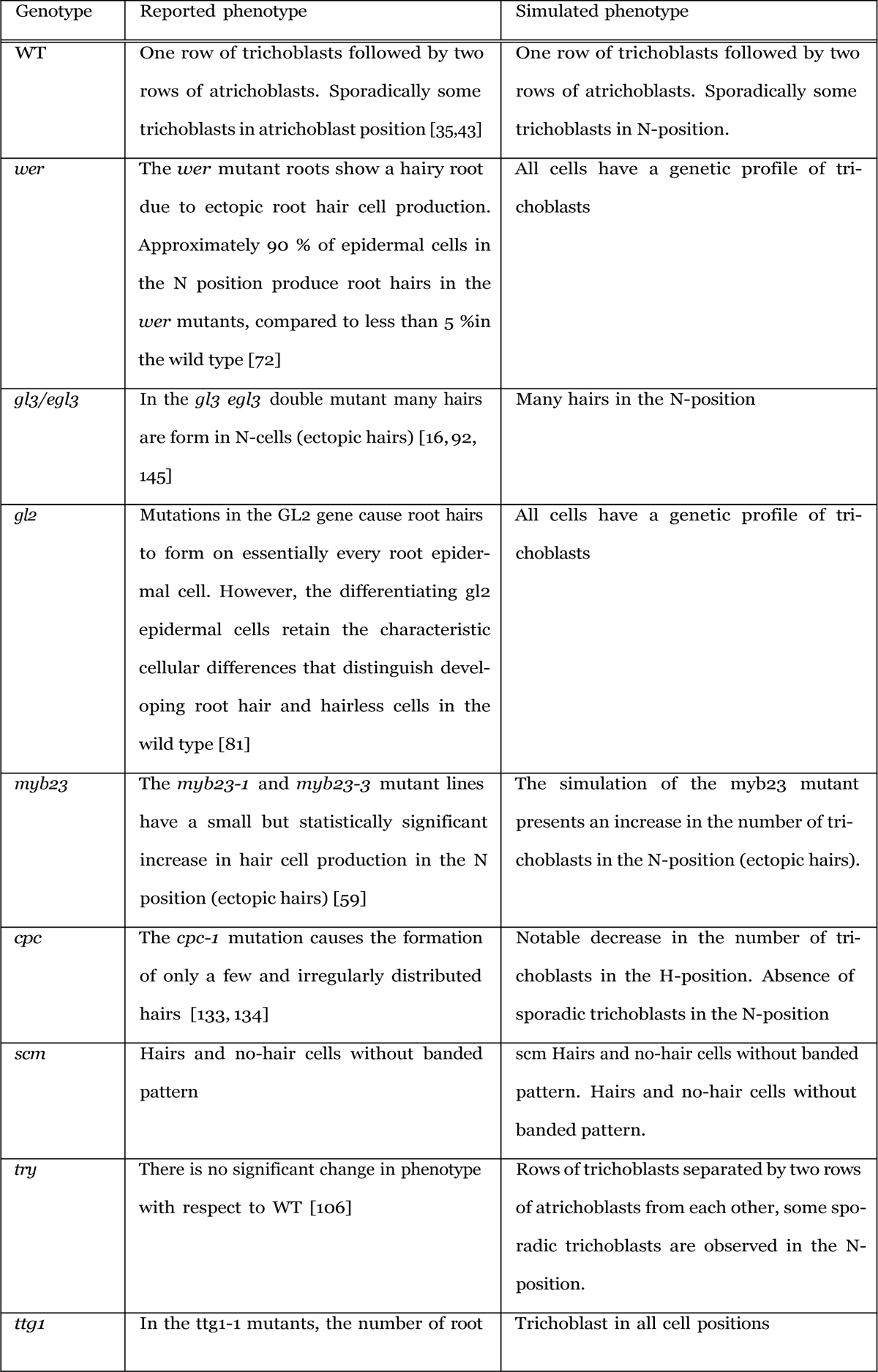
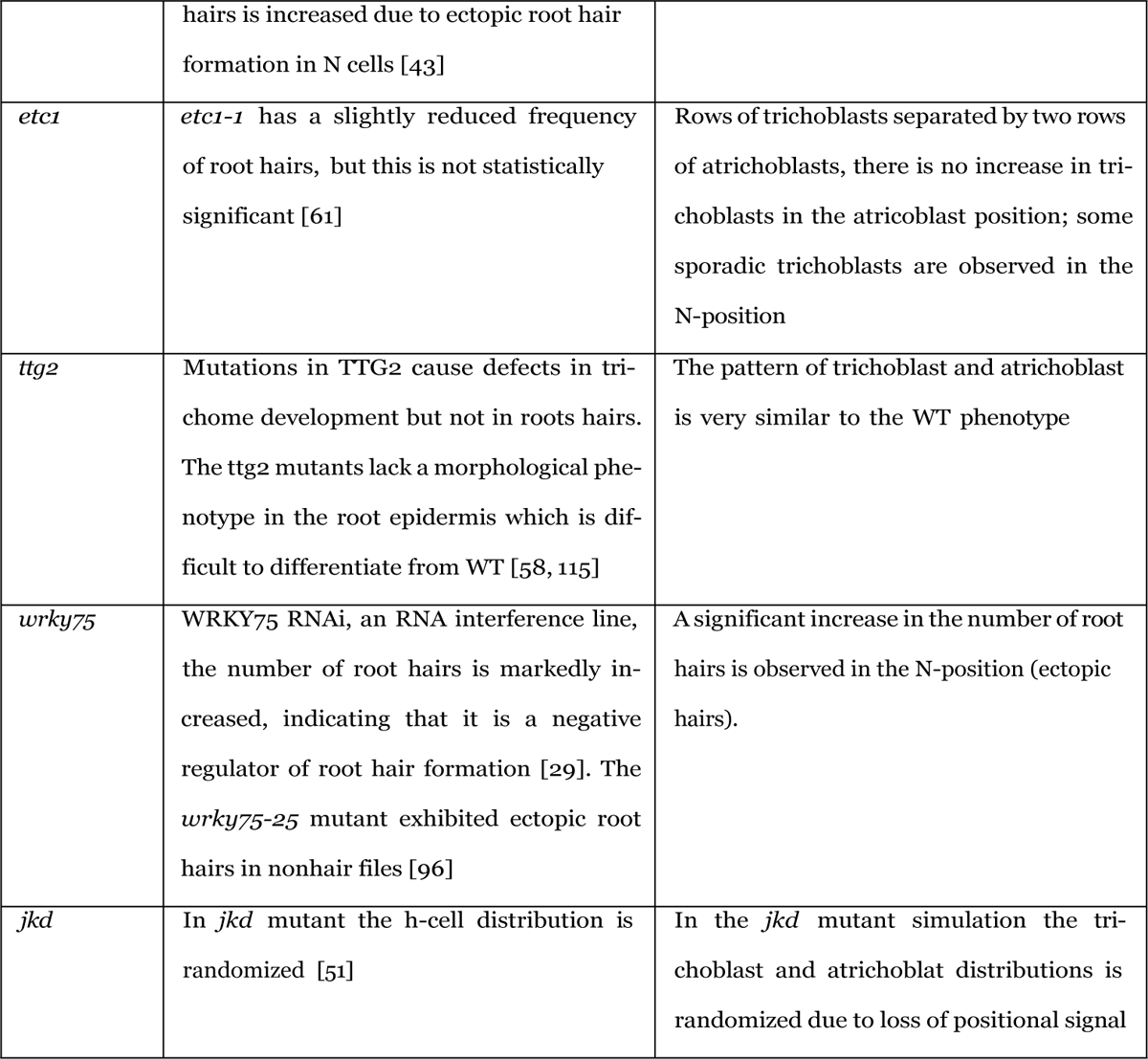
Comparison between reported phenotypes and simulated phenotypes.

**Table 6.**
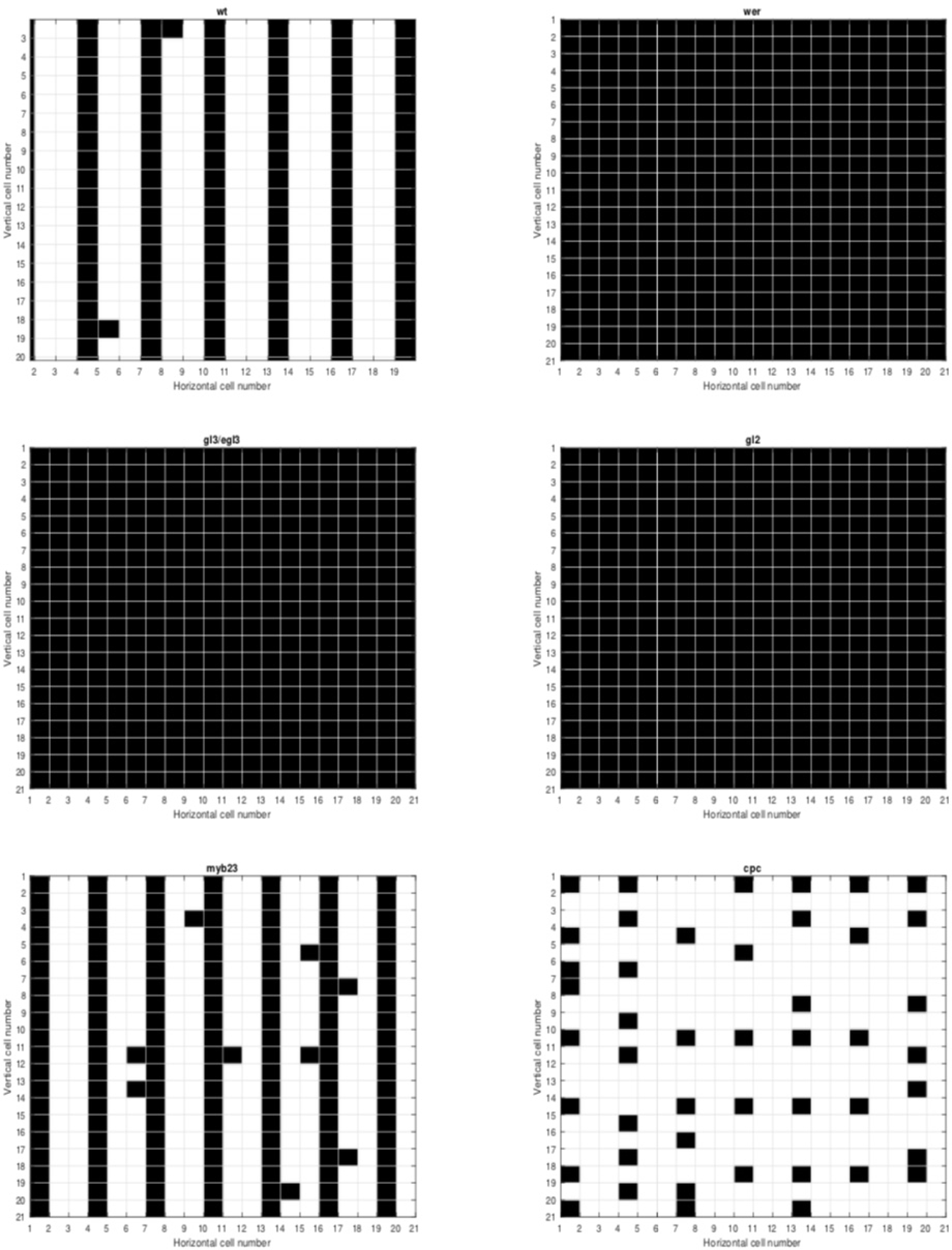
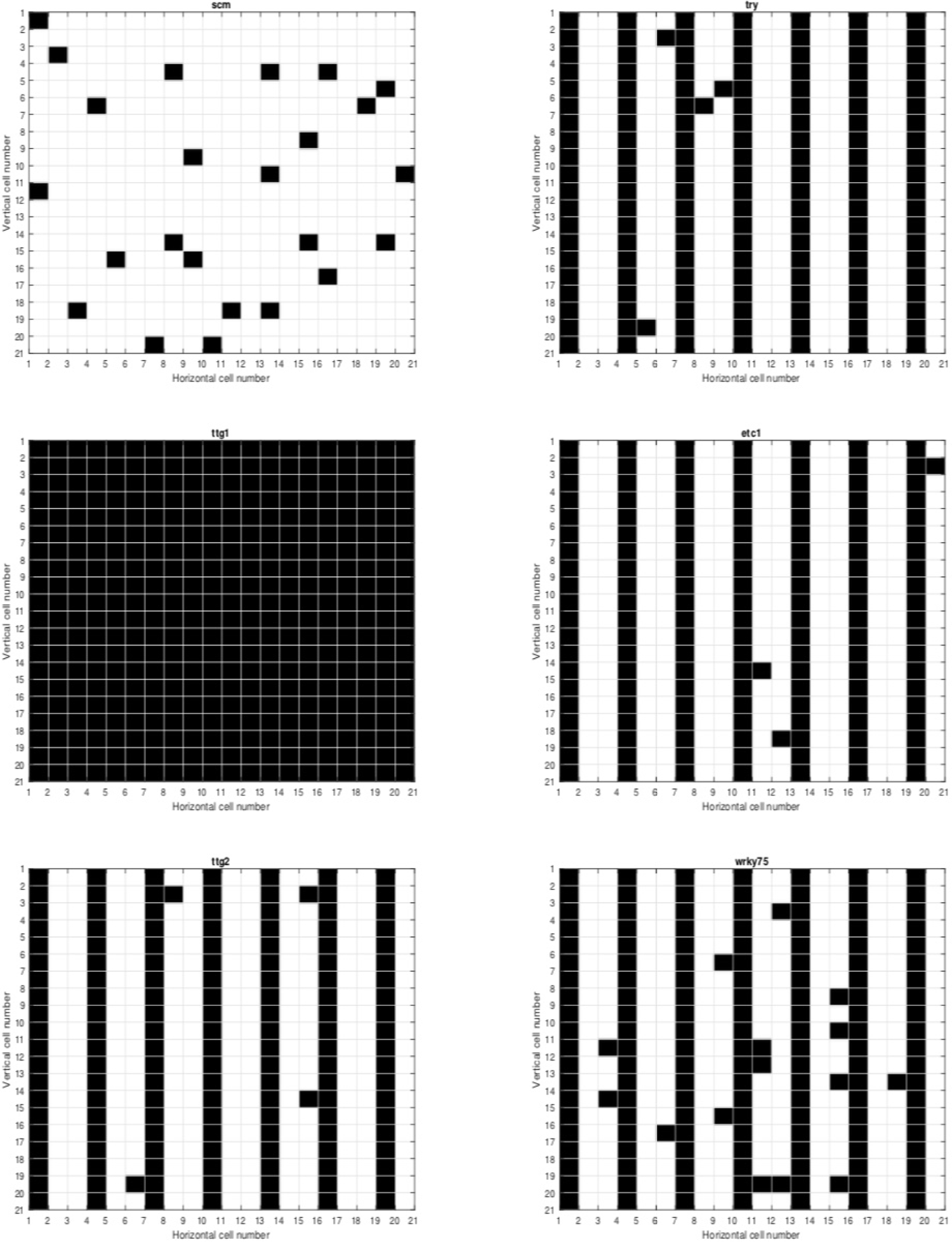
Spatial patterns of the simulated mutants.

**Table 7.**
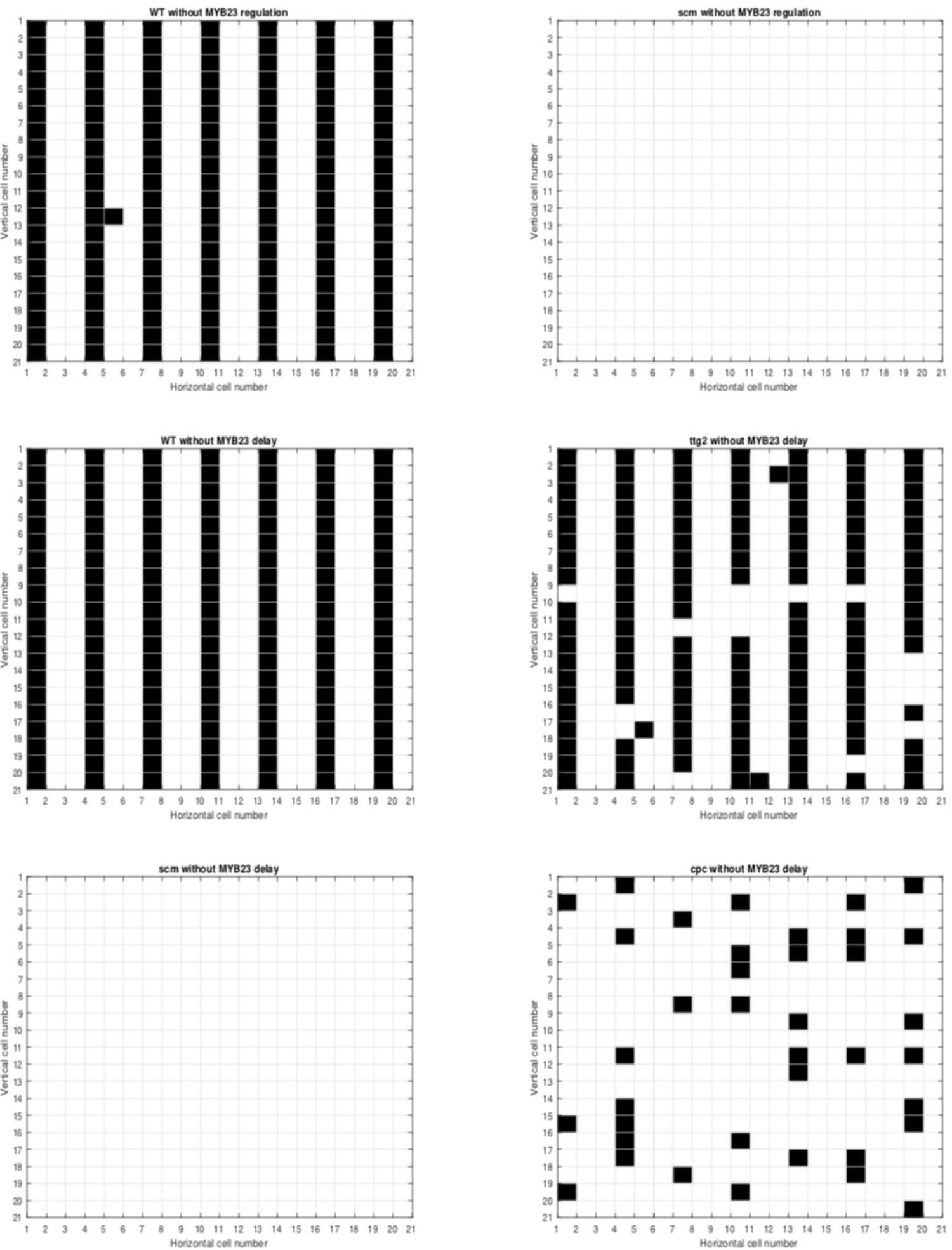
Simulated spatial patterns without the proposed interactions.

**Table 8.**
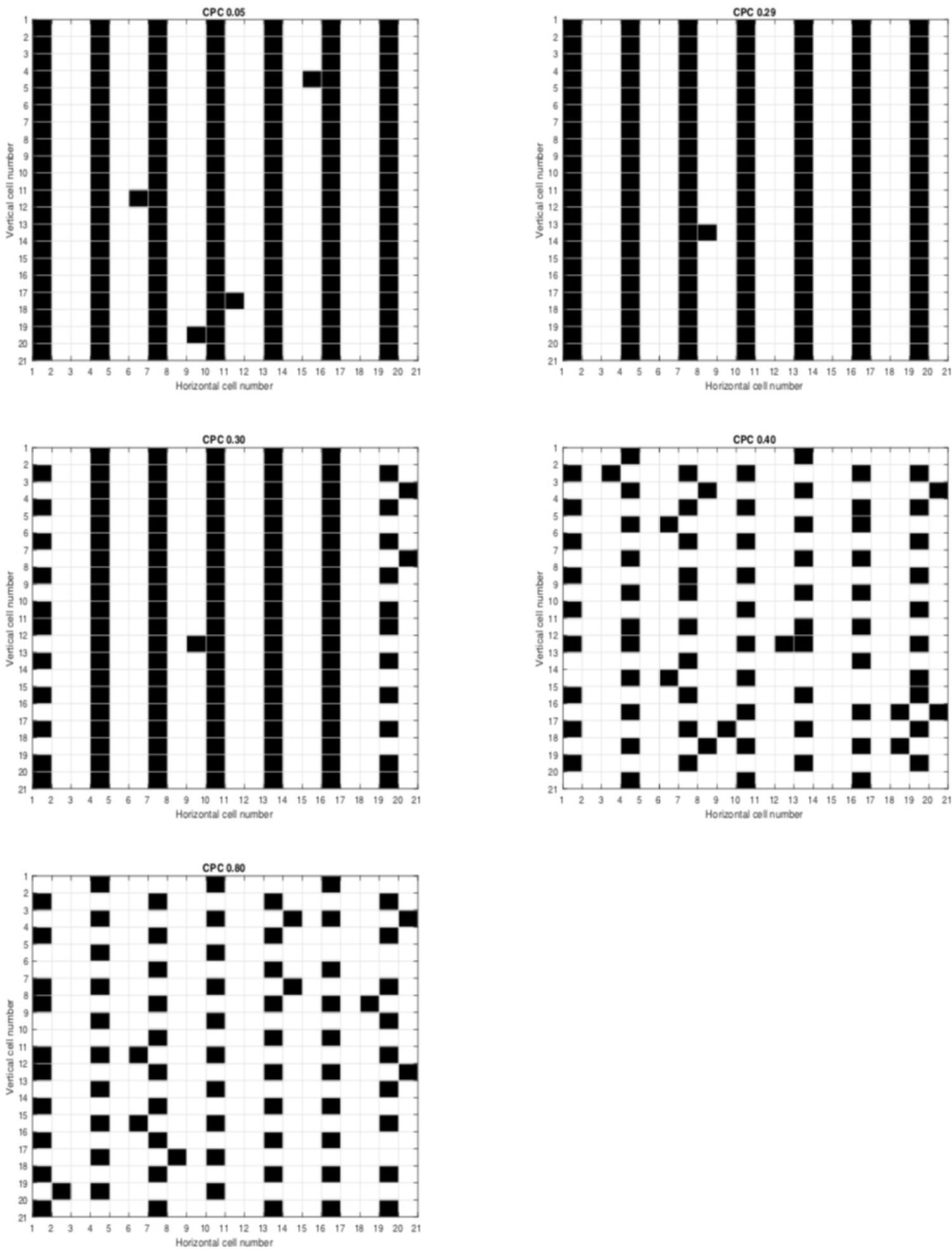
Patterns of organization of trichoblasts and atrichoblasts for different values of diffusion of CPC.

An interesting aspect of our model model is the fact that in this mutant GL2 expression levels are high compared to WT while GL2 protein levels are similar to WT, which could suggest an important post-translational regulation [137]. To explore this hypothesis it is possible to follow two possible approaches using our model: A first approach is the modification of the dynamic rules of WER activity, which, as explained in previous paragraphs, could describe the reduced binding of WER to CPC and TRY, and evaluate the effect on the relative accumulation of the BMW complex. This approach would also involve considering different levels of WER expression that correlate with the ability of the WER D105N protein [137] to bind with low affinity to CPC and TRY. The second approach involves making the relevant modifications in the diffusion parameters of the CPC protein and in the rules that regulate the lateral inhibition mechanisms to allow a bidirectional mobility mechanism of the protein that does not depend on the relative position. In both cases, these modifications will allow us to explain two points of the previous model: the role of the relative accumulation of CPC at the N-position and at the H-position, and the activity of WER, via the MBW complex in the regulation of the non-position-dependent expression of CPC and its accumulation. At the moment this hypothesis seems to make sense in the light of the model proposed here. As can be seen from the network structure, GL2 is located furthest downstream and is targeted by other important regulators. The simulations obtained correspond to those reported in the literature for the *gl2-1* mutant [119]. From the comparison, it can be seen that this phenotype is a result of the increased number of trichoblasts in the N-position (see Table 2 and Table 3). It has been described that even though in this mutant the cells in the N-position have requested their ability to differentiate into atrichoblasts, these cells exhibit a greater length and early vacuolization with respect to cells in the H-position as observed in the wild type [43, 81].

For the *gl2* mutant, simulation results show that all cells differentiate to trichoblast as reported [81]. The comparison between the percentages of trichoblasts in the simulations and the experimental evidence is similar, with a small variation of 0.8% in the number of atrichoblasts in the N-position. [119]. It has been suggested that GL2 is responsible for only part of the atrichoblast identity and that it is the upstream regulators in the GRN that regulate the developmental events of atrichoblasts. Following cell pattern determination, it may be inferred that GL2 promotes a portion of N cell differentiation (differentiation that does not result in the formation of a root hair) and that the upstream trans-activating complex may regulate every aspect of N cell development, including the cell’s fate. The *gl2* mutations are believed to act in a semi-dominant way during N cell differentiation since roots heterozygous for the gl2 mutations *gl2-1* and *gl2-2* develop ectopic root hairs in N cell files with noticeably higher rates than do wild type roots [81]. A small amount of GL2 expression in the H-position is also assumed to be necessary for normal root hair tip growth, in addition to GL2’s high level of expression in N-cells. This is because GL2 mutant root hairs branch out much more frequently than wild type root hairs. [91]. In the context of this model, the possibility of evaluating a possible basal expression of GL2 in H-position cells may be possible by converting the GL2 node into a discrete variable with several levels of expression, however, since our model is focused on the cell determination process and not on the initiation and tip-grow process. On the other hand, it is possible to extend the proposed model by converting the logical variables and transforming it with a fuzzy logical approach to obtain a system of differential equations with which to evaluate changes in GL2 concentrations at both cell positions [31].

In the case of the TTG1 mutant and the double mutant *cpc ttg1*, the results obtained are interesting. As described above, the results of the simulations of the *ttg1* mutant are similar to those reported for the *ttg1-1* mutant, so this model recovers in a consistent way the experimentally previous results for this mutant. In addition to the ttg1-1 allele, several other mutant alleles have been described for the TTG1 gene. These mutants are *ttg1-1* [62], *ttg1-13* [70], *ttg1-9* [43], and more recently the mutant alleles designated *ttg1-23 and ttg1-24* [80] have been described. The identification of these mutants and their very different phenotypes raises a number of questions regarding the results of our model. As described above, for the simulations of the *cpc ttg1* mutant, the most similar phenotype reported in the literature in pattern and proportion of trichoblasts and atrichoblasts corresponds to *cpc-1 ttg1-13* [80], However, only partially, since in our simulations we observed a considerable increase in the number of trichoblasts in the H-position and a lower number of atrichoblasts in the N-position. These results suggest that the double mutant *cpc ttg1* generates a higher number of ectopic hairs in our simulations. Experimental evidence suggests that of the alleles identified so far for TTG1, the *ttg1-13* and *ttg1-1* mutants show the hairiest phenotype with an important percentage of ectopic trichoblasts while *ttg1-9*, *ttg1-23* and *ttg1-24* mutants have a lower percentage of ectopic trichoblasts, demonstrating that very strong mutations in the TTG1 gene negatively regulate the differentiation of atrichoblasts and weaker mutations prevent the differentiation of ectopic trichoblasts [80]. The regulatory implications of this fact in the context of the network are interesting since they could explain the partial recovery of the *cpc ttg1* mutant in our simulations. In general terms, the weak alleles show a GL2 expression similar to WT, whereas the strong phenotypes show reduced GL2 expression, and in the weak alleles the RNA levels of the CPC gene are higher. In all reported alleles TRY levels are decreased, while MYB23 expression levels seem not to be altered and finally TTG1 protein expression patterns are similar for all alleles to those reported in the WT. Together these results explain part of the disjunction of this mutant and how our model could be suitable to recover this and the other phenotypes described for the different TTG1 alleles [80]. As ttg1-13 is a strong allele, the expression levels of GL2, TRY and CPC are decreased with respect to WT and with respect to the weak alleles. Consequently, there is less TRY and CPC, now that they are immersed in the *cpc-1* mutant, the effect of decreased CPC expression would further contribute to lower total CPC/TRY levels, which would contribute to a higher differentiation of atrichoblasts, however, since GL2 expression levels are low, the differentiation of atrichoblasts decreases allowing the differentiation of trichoblasts in the epidermis, which explains why in our simulations the *cpc ttg1* mutant is completely hairy. In order to modify our model and explain the phenotypes of the other alleles such as the double mutants with *cpc,* a possible route is the incorporation of new logical rules that allow that in such mutants the GL2 expression levels are higher even when with respect to WT the levels of CPC and TRY are low but higher than those observed in the strong alleles. And secondly, that the mobility of the CPC protein is higher than in the current model given that the levels of CPC RNA are higher in the weak alleles [80] than in the stronger, which together with the rules would allow a greater amount of the MBW complex and therefore that despite the *cpc* mutant background, the phenotype of the *cpp ttg1* double mutant enhances the CPC phenotype and a basically hairless phenotype is observed.

### The crucial role of intercellular communication and GRN element diffusion in determining the fate of epidermal cells

Lateral inhibition is an important proposed mechanism of pattern formation in multicellular organisms that has been studied extensively for its ability to explain cell determination and patterning mechanisms [23, 83, 117]. Different examples have been described during development in plants whose underlying mechanism is the existence of a lateral inhibition mechanism: In *Marchantia polymorpha*, a miRNA-mediated lateral inhibition controls rhizoid cell patterning [122]. On the other hand, it has been described that the initiation site where the lateral roots of the primary root of *Arabidopsis* will form is controlled by a signaling cascade involving a lateral inhibition mechanism mediated by the peptide hormone TOLS2 [128]. As mentioned, it has been suggested that this same dynamic mechanism is underlying the spatiotemporal pattern of cell fate determination of trichoblasts and atrichoblasts [73, 94, 106, 107, 110]. In this context is the activity of the gene CPC (and the mobility of the protein codified by it) that helps to generate the proper accumulation pattern of the WER-GL3/EGL3-TTG1 complex [108]. Thus the role of CPC in modulating the lateral inhibitory mechanism underlying the GRN structure has been well documented [17, 108].

As can be observed in the results of the analysis of the diffusion parameters of the key transcriptional factors in the determination of the cell types of the epidermis and as has been suggested in the literature, the diffusion and movement of these play an important role in the establishment of the pattern of organization of the epidermis of the root of *Arabidopsis* [104]. The analysis of diffusion parameters of key transcription factors reveals their significant role in determining epidermal cell types, as supported by existing literature. The diffusion and movement of these factors are crucial in establishing the organizational pattern of the *Arabidopsis* root epidermis [104]. However, our results demonstrate that understanding the role of diffusion requires considering the behavioral context of the underlying genetic network that governs the cell determination process between atrichoblasts and trichoblasts [10, 11, 13].

As can be seen from the results of the simulations, the isolated and joint variation of the diffusion values of CPC and GL3/EGL3 modifies the epidermal organization pattern of the WT phenotype, with the pattern being more sensitive to changes in GL3/ELG3, while for CPC the changes are less drastic in the pattern organization (See Figure 3 and Table 6). As discussed, the role of CPC in the establishment of this lateral inhibition mechanism is important to highlight. We have tried not only to elucidate its role within the GRN but also the structural mechanisms of its protein that regulate its cell-to-cell movement capacity. CPC is part of a family of several homologous proteins including ETC1 and TRY [136], but of all of them only CPC is capable of cell-to-cell movement [64, 133]. It has been previously established that the S1 and S2 motifs are necessary in the structure of the CPC protein for its mobilization, being the S2 motif also necessary for its nuclear localization [64]. Thus, and given the structural and functional similarities of this protein family, it has been thought that other proteins homologous to CPC could have the ability to mobilize between cells of the epidermis and establish new feedback loops that allow giving robustness to the GRN, especially due to the conservation of the S2 motif among the various members of the family [108]. However, so far, this hypothesis has not been proven [125].

With the help of chimeric constructs, some research groups have attempted to analyze the effect that the different motifs (S1 y S2) and amino acid residue changes may confer on the mobilization capacity of CPC and its homolog proteins and the root phenotype generated by them. For example, in the chemical protein S1:ETC1, it is not possible to observe its mobilization from non-hair cells to hair cells, suggesting that while in CPC the S1 motif is necessary for mobilization, in the ETC1 protein it is not able to generate this behavior. Detailed sequence analysis of the CPC and ETC1 protein sequence shows important differences in the contiguous S1 motif region, in particular, positions 21, 22, 24, 37, 40, and 75 of their amino acids are very different in charge and polarity, which could confer significant differences in the protein structure that would allow the mobility of CPC and not ETC1. [125]. On the other hand, in a series of subsequent experiments it has been demonstrated that the modification of the CPC protein structure with the incorporation of specific sequences of the ETC1 protein is possible to modify the mobilization capacity of the CPC protein: Of the five chimerical proteins analyzed, four maintain the ability of CPC to mobilize, all of them with changes in point amino acid residues, however two of them (Chimera 1 and Chimera 2) in which the N and T amino acids of ETC1 were introduced between positions 11 and 12 of CPC and the introduction of the HLKTNPTIV sequence of ETC1 at positions 21 and 22 of CPC, respectively, showed lower GFP fluorescence intensities in hair cells compared to non-hair cells, suggesting that although they maintain their ability to mobilize between epidermal cells, they show a temporal delay, *i.e.* a decrease in the rate of mobilization [126]. It is important to note that the roots of the plants expressing these chimeric proteins showed many root hairs for the increase in the number of ectopic hairs, however, how the proportion of trichoblasts and atrichoblasts in the H-position and in the N-position is altered is something to be analyzed in these lines [126].

Previous results of the CPCp::CPC:GFP plants show that their phenotype is similar to that observed in the 35S::CPC overexpression lines in which 84.8% of hairs are observed in the N-position and a decrease to 15.2% of non-hair cells in the N-position [64]. Relating this evidence to our results we can notice two things: firstly, it is possible to modify the patterns of organization of trichoblasts and atrichoblasts by modifying the diffusion capacity of the CPC protein by altering specific amino acid residues in its structure, and secondly, it is possible that the alterations in the N-position of the CPC protein may be due to the alterations in the N-position of the N-position of the cells [64], it is possible that the alterations in the number of total hairs observed in these mutants are the result of altering the trichoblast/atrichoblast ratio at the H-position and and the Nposition, and as a result, the spatiotemporal dynamics of the GRN that underlies the differentiation of these two cell types.

Our results suggest that the increase in CPC diffusion generates a decrease in the number of total epidermal hairs due to the loss in the number of trichoblasts in the H-position, however low values of the Dcpc parameter generate phenotypes similar to those observed in WT. Comparison with what has been previously discussed highlights that if as in the case of CPC chimeras 1 and 2 [126], what is observed is a possible decrease in the diffusion capacity of CPC and therefore a delay in its transport. In that sense, our model gives only partial results to be able to explain these data. Given that the simulation approach proposed here uses a discretized Laplacian that depends on the threshold parameter to be able to transform the diffusion values and thus be able to update the dynamics of the network, possibly part of the behavior of the diffusion process of the CPC protein is being lost, in such a way that for very small values of diffusion, we cannot observe this ectopic increase due only to the decrease in the CPC diffusion rate, as if we can see it for large values where the opposite effect to that reported for the chimeric constructions previously described can be observed. Recent experiments in which another group of three chimeric proteins was analyzed to see the role of key amino acid residues of the CPC protein point in see sense [54]. Of the 3 chimeric proteins generated (Chimeras 6, 7, and 8) all were generated by point changes in positions, it has been described that chimera 8.

### Uncovering the impact of novel interactions and feedback loops on epidermal cell patterning

As discussed in the previous paragraphs, the model is able to recover several mutant lines of loss of function qualitatively and quantitatively by comparing with the results reported in the literature. Also with the model, we are able to explore the role of diffusion of mobile elements between trichoblasts and atrichoblasts in the modeled cell domain and demonstrate the importance but not sufficiency of diffusion in establishing the characteristic pattern of organization of *Arabidopsis thaliana* root hairs.

The results of our simulations indicate that the proposed new interactions within the GRN are capable of recovering most of the phenotypes reported for the genes under consideration in a robust way. These results highlight the significance of feedback loops and diffusion for the appropriate establishment of the epidermal organization pattern. We will go into more detail about the relevance of these novel interactions within the GRN structure in the sections that follow, as well as how they may affect how epidermal cell type determination dynamics operate.

Previous computational models have suggested the central importance of positive and negative feedback loops within the GRN and their dynamic implications for the determination of *Arabidopsis* root epidermal trichoblast and atrichoblast [9, 11, 94, 100]. At the center of these feedback loops is the activity of WER. As previously described, WER is part of the MYB-bHLH-WD complex whose main function is the activation of GL2 and the inhibition of trichoblasts. The function of this complex is diminished by the competitive binding of CPC [134]. We can divide the regulatory mechanisms underlying the GRN structure of this model as follows: a) position-dependent feedback mediated by the activity of JKD and SCM [51, 65], b) positive feedbacks of the MBW complex, c) negative feedbacks on the MBW complex, d) lateral inhibition mechanisms mediated by diffusion of CPC and GL3/EGL3 proteins and e) mechanisms that reinforce the cell pattern. One important regulatory mechanism is the regulation of the activity of SCM by the WER-GL3/EGL3-TTG1 complex. Despite SCM being expressed in all root cells, except the root cap cells [66] its activity is decisive as a positional information mechanism [68]. In the *wer-1* and *gl3 egl3* mutants, the SCM::GUS reporter is more highly expressed in different root tissues compared to the wild type, suggesting that the MBW complex is an important negative regulator of SCM activity. The results of our simulations show that the incorporation of the regulation of the MBW complex on SCM activity is important and the results of the simulations of the *scm* mutant demonstrate that the loss of the positional signal allows the pattern to be altered even when trichoblasts differentiation occurs but in a random manner over the simulated domain. Simulations of the *scm* mutant in which the mechanisms regulating its activity are altered (in any of the regulators of the MBW complex or its elements) show unobserved behavior with a tendency to eliminate trichoblast differentiation and loss of positional signal. The fact that our model only partially recovers the pattern of the *scm* mutant raises the hypothesis about other regulators involved in its regulation, particularly the importance of the regulators JKD, and QUIRKY [118] (See below), and about the regulatory mechanisms that allow sensing of the positional signal relative to the cortex cells.

According to [59], experimental evidence suggests that MYB23 expression takes place during a later stage of root development compared to WER expression, its role in determining the pattern could be more related to the subsequent stabilization of the pattern once WER has begun its activity, establishing a reinforcement of the pattern [59].

When MYB23 expression was allowed from the beginning of the simulations, there was no significant alteration in the characteristic pattern of the WT phenotype’s organization. However, in the *scm* mutants, a notable reduction in the number of trichoblasts was observed. Some simulations did not form any trichoblasts, while others exhibited an average of 5 trichoblasts across the epidermis (refer to Supplementary Information). These findings indicate that trichoblast formation in the epidermis relies on the positive feedback loop of the MBW complex through MYB23 activity, as well as its temporal regulation in relation to WER activity. This process operates independently of the positional signal established by SCM. Finally, the regulation of the MBW complex activity by the TTG2 gene is necessary for the recovery of the *scm*, etc1 and try mutants, as well as for the correct determination of the sporadic ectopic trichoblasts observed in the WT phenotype. This interaction is important since TTG2 appears to form a positive regulatory loop on the activity of the MBW complex in addition to that determined by the MYB23 gene, a fact that is demonstrated by the necessity of this interaction for the correct determination of trichoblasts in the *scm* mutant. Therefore, the positive regulatory loop of TTG2 incorporated in this model on the activity of the MBW complex allows reinforcing the pattern independently of MYB23 and establishes itself as an additional regulatory mechanism on the identity of trichoblasts.

### Other interactions and limitations of the model

The relationship between cell specification of trichoblasts and atrichoblasts and cell-cycle regulation is another crucial aspect to investigate. Research has shown that cell fate specification in root hair patterning is influenced by cell division [15]. A regulatory link between cell-cycle and cell fate specification in the root epidermis has been demonstrated through the GL2 EXPRESSION MODULATOR (GEM). GEM is regulated by CDT1, a component of the DNA pre-replication complex, and negatively regulates the expression of TTG1, dependent on the cell cycle. Additionally, GEM is involved in maintaining the repressive state of histone H3 in the GL2 and CPC promoters [20].

Similarly, it is crucial to explore the potential inclusion of additional regulators that have not been considered in this model. For example, the genes QUIRKY (QKY), TORNADO1 (TRN1), and TORNADO2 (TRN2) play an important role in establishing position-dependent epidermal patterning [22, 130]. TRN1, a protein with a leucine-rich repeat ribonuclease inhibitor-like domain, has been found to be essential for ensuring the correct formation of epidermal patterns [69]. In the *trn1-t* mutant lines the GL2 expression pattern is very similar to that observed in the *smc-2* mutant, WER expression is observed in almost all epidermal cells and finally, EGL3 expression is scrambled. This suggests that TRN1 is important in the regulation of preferential expression of WER in N-position cells and its alteration generates ectopic expression of WER in the root leading to a disorganization of the epidermis by altered expression of EGL3 and GL2 [69]. On the other hand, the double mutant *cpc trn1-1* has a lower number of hairs than those observed in the *cpc* mutant and it has been suggested that this effect is explained by the additive effect of the alteration of WER expression and the loss of CPC function. Furthermore, analysis of the double mutant *ttg1 tnr1-1* suggests that *ttg1* is epistatic to *trn1*, which could be supported by the fact that TRN1 regulates WER, and WER in turn regulates the integrity of the MBW complex. The exact mechanism by which TRN1 regulates WER expression is not yet clear, nor is it clear whether or not this mechanism is similar to that observed in other WER regulators, particularly SCM; perhaps it could be part of a parallel feedback loop that stabilizes WER activity [22, 69].

In parallel, QKY, a protein with multiple C2 domains and a transmembrane region (MCTP), has been previously documented to engage with SCM, thereby influencing the patterning of epidermal cells [130].QKY functions as a regulator positioned upstream of the GRN and acts over WER and GL2 functions. Experimental evidence suggests that QKY plays a pivotal role in enhancing the cell-to-cell movement of CPC between epidermal cells by stabilizing SCM protein at the plasma membrane. Furthermore, the experimental evidence demonstrates a direct physical interaction between QKY and SCM via their extracellular domains. Moreover, QKY prevents degradation of SCM by preventing its ubiquitination [118]. Particularly this interaction could be of importance in the dynamics of *cpc, scm wer*, *scm cpc* mutants. In such simulated mutants, our simulations recover in general terms the organization pattern but not the proportion of trichoblasts and atrichoblasts reported in the literature at the H-position and at the N-positions. For the *scm* mutant, our results show a lower proportion of trichoblasts and a higher proportion of atrichoblasts, indicating that the recovered phenotype is more drastic than the one reported for this mutant.

Given the importance of QKY in the stabilization of SCM and its participation in the preferential accumulation of SCM at the H-position as well as its importance in the selective accumulation of CPC at the H-position and thus the inhibition of the MBW complex via WER inhibition, this reinforces the pattern and allows for proper trichoblast differentiation [118]. In such a case, the incorporation of QKY into the model could establish a positive feedback loop on the activity of the inactivation complex and therefore in mutants in which the model recovers a lower percentage of trichoblasts in the H-position, perhaps the pattern is less drastic and the number of trichoblasts in H-position is slightly higher.

In the case of the mutants whose simulations recover a higher percentage of trichoblasts and therefore a lower percentage of atrichoblasts, this could result from the fact that in the model WER activity is regulated by two pathways: the positive activity of SCM and the inhibition of the inactivator complex. In such a case, the incorporation of QKY could be determinant in the regulation of WER activity since QKY facilitates CPC mobilization while it would be a reinforcing mechanism of the negative regulatory loop on MBW complex activity via WER inhibition and therefore could be an additional mechanism to the inhibition of trichoblast differentiation activity, a possible evaluation of these interactions could be necessary in a much more detailed model.

One of the feedback mechanisms responsible for maintaining the stability of the root hair organization pattern involves the action of MYB23. As previously mentioned, MYB23 functions redundantly with WER [59]. Experimental evidence indicates that in the pum23-4 mutant of the ARABIDOPSIS PUMILIO23 (APUM23) gene, which encodes a Pumilio/PUF domain protein primarily associated with impaired ribosome biogenesis, the expression of MYB23 increases through a distinct pathway. Importantly, this pathway is not reliant on the activation of the MBW complex but necessitates the involvement of the ANAC082 gene, a member of the NAC family. The results of this increased expression is the ectopic non-hair cell production of the root epidermis [138]. The importance of this mechanism is not clear, however, the existence of non-canonical regulators within GRNs that underlie epidermal cell type differentiation could be related to the mechanisms of phenotypic plasticity of the epidermis in response to various plant stresses [138]. The contextual significance of the canonical mechanisms observed in the WT and reported in this work seems unclear, but their exploration through approaches such as the one proposed here could help to elucidate the role of these parallel pathways within the GRN and their relationship to normal and altered root epidermal morphogenesis.

An important aspect to consider is that our modeling approach has certain limitations when modeling the activity of the different elements of the GRN, particularly in those where there are expression levels that cannot be expressed as discrete values as we have proposed in this model. As can be seen in some of the mutants whose phenotypes could only be partially recovered in the simulations because our simulation approach is unable to consider subtle variations in the expression patterns of the different regulators within the GRN. In the case of mutants whose patterns and proportions of trichoblasts and atrichoblasts could only be partially recovered, the double mutant coc myb 23 is found. Neither is it possible to finely recover the variation in the accumulation of the activator and inhibitor complexes as has been reported. However, it could be considered an extension of our model. This takes into account the following considerations: first, it is possible to propose a modification of the model in order to reduce it to a Boolean model by means of some of the methodologies that have been proposed for this purpose [39, 127]. Once the Boolean GRN, the equivalent network must be analyzed to verify its consistency, that is, that the attractors are equivalent to those obtained in the discrete model, so that the expression profiles of the stationary states correspond with the expected attractors defined by experimental data. It is also necessary to analyze aspects such as the robustness of the model and finally the simulation of gain and loss of function mutations [25]. Once the equivalent network is coherent with the results presented here, it is then possible to propose a system of ordinary differential equations using the logical rules, for which multiple methodologies have been proposed [4, 86, 132]. The resulting continuous system can then be analyzed to explore finer behaviors such as the timing of the interactions or modeling concentrations of transcriptional factors to explore their relative abundance. On the other hand, it is possible to carry out bifurcation analysis in order to explore the propensity of certain nodes to allow transit between attractors by modifying parameters of the functions that determine their dynamics [26]. Finally, with the resulting system of equations, it is possible, in the context of a meta-GRN, to propose a spatial model using a methodology similar to the one proposed here or to use others such as the Cellular Potts model (CPM) in which it is possible to model in a way more final the processes of diffusion and transport [121]. In our group, we have already made approximations in this sense to explain the emergence of the organization patterns of the root niche of *Arabidopsis* [45].

### Perspectives and future directions

The type of approximation described here also offers a valuable opportunity to investigate the phenotypic plasticity of the *Arabidopsis* root epidermis. Research has shown that trichoblasts and atrichoblasts differentiation and their organizational patterns, respond to changes in environmental conditions, especially nutrient availability [89, 101]. It has been observed that the differentiation of cell types in the root epidermis responds differently to deficiencies in various elements, such as phosphates [21, 30, 57], cadmium [5, 6], arsenic [5] nitrates [78], salinity [139], and CO2 [90], indicating a potential avenue of exploration for this approach. In this regard, the use of simulation tools to explore the dynamic aspects of phenotypic plasticity has begun to attract attention in the context of the management of crop species such as corn [42]. From our perspective, the use of cellular-scale multilevel models of the GRN underlying the determination of trichoblasts and atrichoblasts and their response to environmental stimuli could be an important tool to describe the systems-level processes of the plastic response of these regulatory modules in different developmental contexts [44].

Recent studies suggest that the structure of the GRN appears to be conserved in many plant species that show the type III organization pattern of the root epidermis. This genetic network is the result of a process of evolution of regulatory mechanisms, particularly position-dependent ones, from an ancestral network whose spatiotemporal dynamics give rise to patterns very similar to those observed in plants expressing the Type I pattern of root epidermal organization where all cells of the root epidermis are capable of originating a hair. Thus it is the emergence of regulatory mechanisms via regulatory loops that determine gene expression differentially by position relative to the cortex cells that gave rise to an ancestral GRN that gave rise to the molecular mechanisms of the Type III pattern as observed in *Arabidopsis*. This network is consequently the result of evolutionary processes of dynamic mechanisms in which the fundamental role of positional regulation evolved through the establishment of feedback loops that regulate the activity of the MBW complex [148]. In this sense, an important aspect to understand in much greater detail the spatiotemporal aspects of the regulation and evolution of this network can be solved by taking this GRN model as a basis and extending it through two essential aspects: the integration of this network with the information obtained through single-cell rna-seq studies and secondly, its extension in multilevel modeling platforms that incorporate aspects not considered in our modeling proposal.

On the first point, there are several recent approaches that have described the expression profiles of root cells and have tried to establish the developmental trajectories of the different cell types at a single-cell resolution not only for WT but also for some mutants and ambient conditions [19,28,36,79,98,111,147] and some of these approaches even take advantage of machine-learning methods to generate better quality data to identify cell-type marker genes from scRNA-seq data [143]. This approach would allow us to extend our GRN with key regulators that were not incorporated in our model by reconstructing regulatory networks based on massive data [142, 145] and to compare previously proposed conserved loops and incorporate new ones that are able to describe the dynamic behavior of epidermal differentiation in different developmental contexts. These new interactions can be validated in the context of our model to assess their relevance.

On the second point, much has been discussed about the relevance of different multiscale simulation approaches to describe tissue novel pattern formation processes and the different computational platforms on which they can be implemented depending on the objectives of the model [33]. Undoubtedly, all these platforms and approaches have limitations and are developed for specific purposes [40]. Of interest for this model is the Epilog platform that allows the logical simulation of genetic and signaling processes and the coupling of these in a cellular domain using hexagonal grids [131]; this approach has been applied to explain the emergence of dorsal-ventral patterns of organization in sea urchin embryo [41], which shows that approaches such as the one proposed in this work can be easily extended to reduce their limitations. On the other hand, specific platforms have been applied or developed to explore the coupling of intracellular processes with cellular dynamics to understand tissue and organ formation in plants, for example: VirtualLeaf [3, 71, 87], Virtual Plant Tissue [27], CellModeller [38], CellZilla [112], and CompuCell3D [45]. These platforms allow the incorporation of intracellular dynamics such as GRNs based on different formalisms (from Boolean networks to nonlinear differential equations) with cell-cell signaling dynamics through diffusion or active transport processes in cellular domains of very different geometries, which could allow us to solve the limitations previously discussed in our model. In such a case we believe that one way to address the limitations of this model, particularly those related to those mutants partially recovered by our approach, is to extend this GRN through a spatial lattice-based formalism, which shall allows us to extend this model to a continuum approach and where we can simulate the processes of cell diffusion and communication in a more fine-grained manner. It has been demonstrated the utility of this approach in previous models to explain the system-level dynamics of root apical meristem [45, 46].

## Author Contributions

Conceptualization: ACJ ERAB. Investigation: ACJ. Methodology: ACJ ERAB. Resources: ACJ ERAB. Supervision: JCMG ERAB. Writing – original draft: ACJ JCMG ERAB.

## Acknowledgments

This work was presented as a partial fulfillment toward AC-J’s doctoral degree in the“Programa de Doctorado en Ciencias Biomédicas” at the Universidad Nacional Autónoma de México (UNAM). ACJ received fellowship 560839 from CONAHCYT.

## Supplementary material

